# Decoding stage-specific functions of PRC2 uncovers a cell-autonomous role in preimplantation development and a non-cell-autonomous role in primordial germ cell fate

**DOI:** 10.64898/2026.01.20.700679

**Authors:** Chengjie Zhou, Meng Wang, Zhiyuan Chen, Yi Zhang

**Author notes:** These authors contributed equally to this work (Chengjie Zhou, Meng Wang).

## Abstract

Early embryogenesis is accompanied by dynamic epigenetic modifications. While such dynamics are important in cell intrinsic regulation of gene expression, their extrinsic roles in mediating intercellular communication during early embryogenesis is less understood. Using the dTAG system, here we reveal previously underappreciated stage-specific functions of PRC2 in regulating pre-implantation and primordial germ cell (PGC) development. We demonstrate that PRC2 plays important roles in regulating maternal to zygotic transition (MZT), and epiblast (EPI) formation. By systematically analyzing H3K27me3 and H3K4me3 dynamics, we redefine the timing of bivalency establishment and uncover a stepwise mechanism governing bivalency acquisition in early embryogenesis. Moreover, PRC2 regulates proper PGC numbers in the EPI by controlling *Esrrb* expression in the extraembryonic ectoderm (ExE). Thus, our study uncovers a previously unknown cell-autonomous function of PRC2 in preimplantation development and its non-cell-autonomous impact in PGC number regulation, both through interplays between epigenetic-epigenetic and epigenetic-TFs networks.

## Introduction

Following fertilization, embryonic development initially relies on the maternal products inherited from eggs and gradually becomes dependent on the newly synthesized zygotic products ^1^. In the mouse, awakening of the zygotic genome takes places in two waves. The minor wave starts at late 1-cell stage with hundreds of genes transcribed, whereas the major wave starts at late 2-cell stage with thousands of genes activated ^2–4^. The maternal RNA decay and zygotic genome activation (ZGA) together are known as maternal-to-zygotic transition (MZT) ^1, 5^. During MZT, the embryonic genome is also subject to extensive epigenetic reprogramming to erase germ cell signatures and to acquire totipotency. After a few rounds of cell cleavages, embryonic cells gradually lose totipotency and undergo the first cell fate specification at embryonic day (E) 3.5 to form inner cell mass (ICM) and trophectoderm (TE) lineages ^6–9^. During the second lineage specification at E4.5, the ICM cells become epiblast (EPI), which marks the establishment of naïve pluripotency ^10^, and primitive endoderm (PrE). Following implantation, the EPI exits from naïve pluripotency and acquires the formative pluripotency at E5.5 and transforms to the primed pluripotency at E6.5 ^11^. Meanwhile, the specification of primordial germ cells (PGCs) from EPI occurs as early as E6.25^12^.

Polycomb group (PcG) proteins play a fundamental role in mediating epigenetic silencing in multiple biological processes ^13–16^. The Polycomb Repressive Complex 2 (PRC2) catalyzes mono-, di-, and tri-methylation at lysine 27 of histone 3 (H3K27me1/2/3)^17–20^. Essential roles of PRC2 in embryonic development have been previously established by functional studies. Zygotic knockout (KO) of the core subunits of PRC2 (*e.g., Eed, Ezh2,* and *Suz12*) indicate that PRC2 are essential for proper gastrulation but largely dispensable for preimplantation development ^21–23^. Loss of maternal PRC2 has a milder effect on postnatal oogenesis despite having some effects on oocyte transcriptome ^24^, but mainly affects non-canonical imprinting, imprinted XCI, and post-implantation development ^25–31^. While the maternal and zygotic KOs of PRC2 have provided important insights into PRC2’s functions in germline and early post-implantation development, neither approach can capture the initial and direct effect caused by PRC2 loss in pre- and post-implantation embryos. On one hand, the zygotic KO may underestimate Polycomb function before embryo implantation due to maternal compensations. On the other hand, for the conditional KOs starting at early stage of oogenesis, potential zygotic compensatory and/or secondary effects may accumulate overtime and complicate interpretations of phenotypes detected in preimplantation embryos.

To address these technical limitations, here we applied the protein degradation tag (dTAG) system ^32, 33^ to systematically evaluate the PRC2 function throughout mouse pre- and early post-implantation development. We showed that the dTAG system enables rapid and efficient depletion of both EED and H3K27me3 in pre- and early post-implantation embryos. Through stage-specific PRC2-depletion studies, we reveal previously underappreciated functions of PRC2 in regulating maternal-to-zygotic transition (MZT), the first and second cell lineage specifications, as well as PGC development. Our results indicate that PRC2 is required for normal MZT and is essential for ensuring proper EPI formation by preventing premature activation of later-stage developmental genes at E4.5 ICM. In addition, we show that the H3K27me3 and H3K4me3 bivalency is established at E4.5 ICM *in vivo*, which is earlier than previously thought of its establishment after implantation ^34^. Importantly, we demonstrate that PRC2 regulates *Bmp4* expression through controlling the expression of *Esrrb* in extraembryonic ectoderm (ExE), which regulates proper PGC numbers through a crosstalk between ExE signaling to EPI cells. These results demonstrate that PRC2 exhibits cell-autonomous and non-cell-autonomous functions through dual interplay with epigenetic and transcription factors during early embryonic development.

## Results

### Targeted protein degradation reveals that PRC2 regulates preimplantation development

A comprehensive analysis of the expression patterns of PRC2 components revealed that most of them are expressed throughout pre- and peri-implantation development (**Fig. 1a**). Similarly, an analysis of our previous low-input ribosome profiling dataset also indicated that these mRNAs are translated in preimplantation embryos **(Extended Data Fig. 1a**) ^35^. Consistently, immunostaining analyses detected H3K27me3 and the core subunit EED from early 1-cell (before ZGA) to blastocyst stage (**Fig. 1b**). The EED and H3K27me3 signals increase at 2-cell stage (when ZGA occurs) and exhibit higher intensity in the ICM than the TE at blastocyst stage (**Fig. 1b and Extended Data Fig. 1b, c**). This 2-cell increase and ICM-biased enrichment of EED and H3K27me3 suggest a potential role of PRC2 in ZGA and lineage specification during mouse preimplantation development.

**Figure 1.**
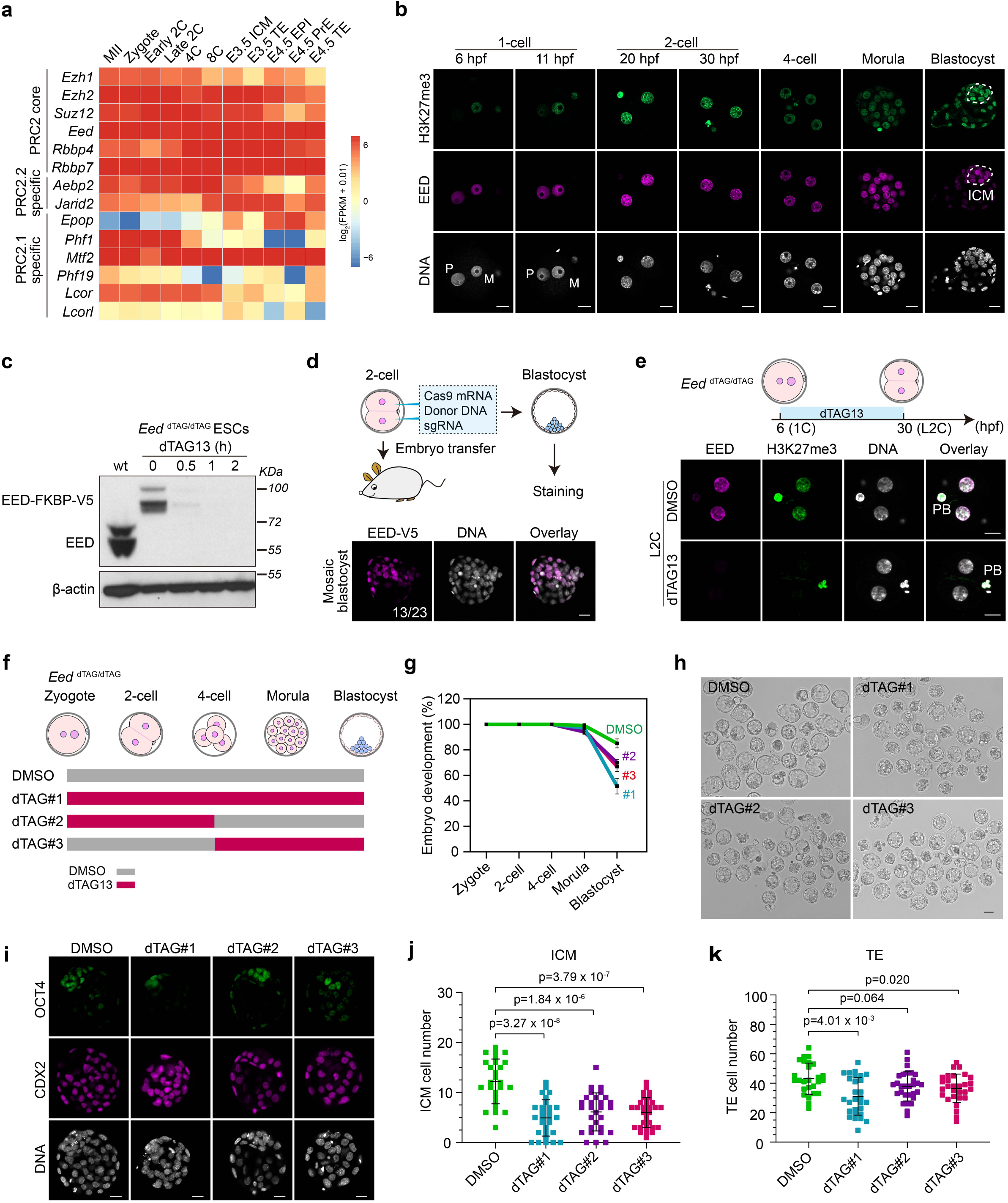
Targeted protein degradation reveals that PRC2 regulates preimplantation development. (**a**) RNA expression dynamics of PRC2 core and accessory subunits during pre- and peri-implantation development. MII: metaphase II eggs; 2C: 2-cell; 4C: 4-cell; 8C: 8-cell; ICM: inner cell mass; TE: trophectoderm; EPI: epiblast; PrE: primitive endoderm; E: embryonic day; FPKM: fragments per kilobase per million mapped reads. (**b**) Immunostaining images showing the dynamics of EED and H3K27me3 during preimplantation development. M: maternal pronuclei; P: paternal pronuclei; hpf: hours post fertilization. The dashed circles indicate the ICM. DNA, Hoechst 33342. Scale bar, 20 μm. (**c**) Immunoblotting images showing the rapid degradation of EED-dTAG protein after dTAG13 treatment in mouse embryonic stem cells. The blots were incubated with anti-EED. β-Actin was used as a loading control. (**d**) Generation of the EED*-*dTAG mouse model. Top panel: schematic of the two-cell homologous recombination (2C-HR)-CRISPR method; bottom panel: immunostaining of V5 tag of the injected embryos at blastocyst stage. The number of injected embryos showing positive V5 signals were shown. DNA, Hoechst 33342. Scale bar, 20 μm. (**e**) Immunostaining images showing the rapid depletion of EED and H3K27me3 in late 2-cell embryos after dTAG13 treatment starting from zygotes. PB, polar body. DNA, Hoechst 33342. Scale bar, 20 μm. (**f**) Schematic showing the dTAG13 treatment strategies during preimplantation development. (**g**) Percentage of embryos that reach the indicated developmental stages in DMSO and dTAG groups (DMSO, *n* = 84; #1, *n* = 69; #2, *n* = 84; #3, *n* = 73; *n* represents the total embryos from three independent experiments). The 2-cell, 4-cell, morula, and blastocyst were evaluated at 28, 48, 72, and 96 hrs post fertilization, respectively. The data are presented as mean values ± SD. (**h**) Bright field images showing the embryos in DMSO and dTAG groups at blastocyst stage. (**i**) Immunostaining images showing the ICM (OCT4^+^/ CDX2^-^) and TE (CDX2^+^) cells at blastocyst stage in DMSO and dTAG groups. DNA, Hoechst 33342. Scale bar, 20 μm. (**j-k**) Quantifications of ICM (panel j) and TE (panel k) cells of panel i (DMSO, n=26; #1, n=28; #2, n=29; #3, n=29). n represent the embryo numbers. *p* values were calculated with Student’s *t*-test (two-sided). The data are presented as mean values ± SD. Experiments were repeated three times. Source numerical data and unprocessed blots are available in source data.

To systematically address the function of PRC2 at different stages of early embryogenesis, we established an *Eed* dTAG model (**Extended Data Fig. 1d**). In the presence of dTAG13, EED-FKBP12^F36V^-V5 fusion protein, is expected to be rapidly degraded by proteasomes. We first confirmed in mouse embryonic stem cells (mESCs) that the EED-dTAG is stable without dTAG13 but undergoes rapid degradation in 30 mins upon dTAG13 treatment (**Fig. 1c and Extended Data Fig. 1e**). We further established a mouse knock-in founder line by co-injecting Cas9 mRNA, the sgRNA, and the donor DNA into the cytoplasm of 2-cell embryos (**Fig. 1d**) as microinjection at 2-cell enhances knock-in efficiency ^36^. Indeed, a high percentage of blastocysts (56.5%, 13 out of 23) showed mosaic V5 tag signals (**Fig. 1d**), suggesting successful knock-in. Importantly, the *Eed* dTAG homozygous knock-in mice, referred to as *Eed^dTAG/dTAG^*, are viable and fertile (**Extended Data Fig. 1f**), suggesting that the EED-dTAG fusion protein function equivalently as the wild type EED.

To evaluate the depletion efficiency in embryos, *Eed^dTAG/dTAG^* embryos were cultured *ex vivo* with or without dTAG13 from early 1-cell (6 hpf, 6 hr post fertilization) to early (20 hpf) or late 2-cell (30 hpf) stages. Immunostaining analyses showed that EED were near completely depleted by early and late 2-cell stages (**Fig. 1e and Extended Data Fig. 1g-i**), indicating a rapid and efficient degradation. Notably, H3K27me3 was also nearly undetectable upon dTAG13 treatment (**Fig. 1e and Extended Data Fig. 1g-i**). These data indicate that we have successfully established a dTAG mouse model that enables rapid and efficient depletion of EED and H3K27me3 in preimplantation embryos.

We next sought to determine whether rapid EED depletion at different stages impact preimplantation development. Specifically, *Eed^dTAG/dTAG^* embryos were cultured with dTAG13 from zygotes to blastocyst stage, from zygotes to 4-cell stage, or from 4-cell to blastocyst stage (**Fig. 1f**). Remarkably, all treatment groups impaired blastocyst formation with the first group showing the strongest effect (∼33% blastocyst rate decrease) (**Fig. 1g, h**). This defect is more severe than the previously reported *Eed* maternal KO that caused post-implantation defects ^27, 29^. The more server phenotype could be due to the simultaneous depletion of both maternal and zygotic EED in the current study. Furthermore, *Eed^dTAG/dTAG^*embryos cultured with dTAG13, irrespective of different treatment windows, showed reduced ICM (OCT4^+^/CDX2^-^) and TE (CDX2^+^) cells (**Fig. 1i-k**). The dTAG treatment caused blastocyst defects in *Eed^dTAG/dTAG^*embryos are specifically caused by EED degradation as the same treatment in WT embryos did not affect the ICM or TE cells numbers (**Extended data Fig. 1j, k**).

Since previous studies have shown that zygotic *Eed* knockout does not affect blastocyst formation and only causes developmental failure at gastrulation ^37^, we next sought to directly compare how degron-induced EED protein degradation and zygotic *Eed* KO may differentially affect embryos develop beyond blastocyst stage. To this end, we utilized a well-established pre- to post-implantation 3D *in vitro* culture system ^38^ and found that *Eed*-KO embryos showed similar E5.5 and E6.5 embryo rate compared to that of the DMSO group (**Extended Data Fig. 1l, m**). In contrast, dTAG-treated embryos failed to develop post-implantation structures (**Extended Data Fig. 1l, m**). These results indicate that acute removal of both maternal and zygotic EED leads to an earlier and more severe defect than zygotic *Eed* knockout. Collectively, these data support a previously unappreciated role of PRC2 in preimplantation development.

### PRC2 regulates MZT and represses developmental genes during ZGA

Given that EED loss from zygote to 4-cell stage impaired blastocyst formation (**Fig. 1g-k**), PRC2 might function during maternal-to-zygotic transition (MZT). To determine how EED loss might directly affect MZT, we performed a time-course experiment analyzing the dynamics of EED and H3K27me3 depletion by adding dTAG13 beginning at the early 2-cell stage (18 hpf). Results show that EED is rapidly degraded within 4 hours when H3K27me3 levels are largely maintained at this time point. The majority of H3K27me3 is depleted by 8 hours and completely lost by 12 hours. These findings indicate that although EED can be efficiently removed within 4 hours, complete depletion of H3K27me3 requires an additional ∼8 hours (**Extended Data Fig. 2a**). Consistently, only minor transcriptional changes were observed, with a small number of genes (15 genes) showing upregulation at 12 hours of dTAG13 treatment (**Extended Data Fig. 2b**), suggesting that short-term EED loss is insufficient to derepress transcription.

We next performed RNA-seq on *Eed^dTAG/dTAG^* early and late 2-cell embryos following EED degradation from 1-cell stage (**Fig. 2a, b and Extended Data Fig. 2c, d**). RNA-seq indicated that acute depletion of EED had minimal impact on transcriptome at the early 2-cell stage (**Fig. 2a**), but it caused 350 and 244 genes up- and down-regulated, respectively, at the late 2-cell stage (**Fig. 2b and Supplementary Table 1, 2**). The up- and down-regulated genes are enriched for maternal genes and ZGA genes, respectively (**Fig. 2c-e and Supplementary Table 1, 2**), suggesting a MZT defect. However, acute loss of EED had little effect on the expression of transposable elements (**Extended Data Fig. 2e**). Collectively, these data suggest that EED acute depletion does not affect minor ZGA but causes defects in major ZGA and maternal decay at the late 2-cell stage.

**Figure 2.**
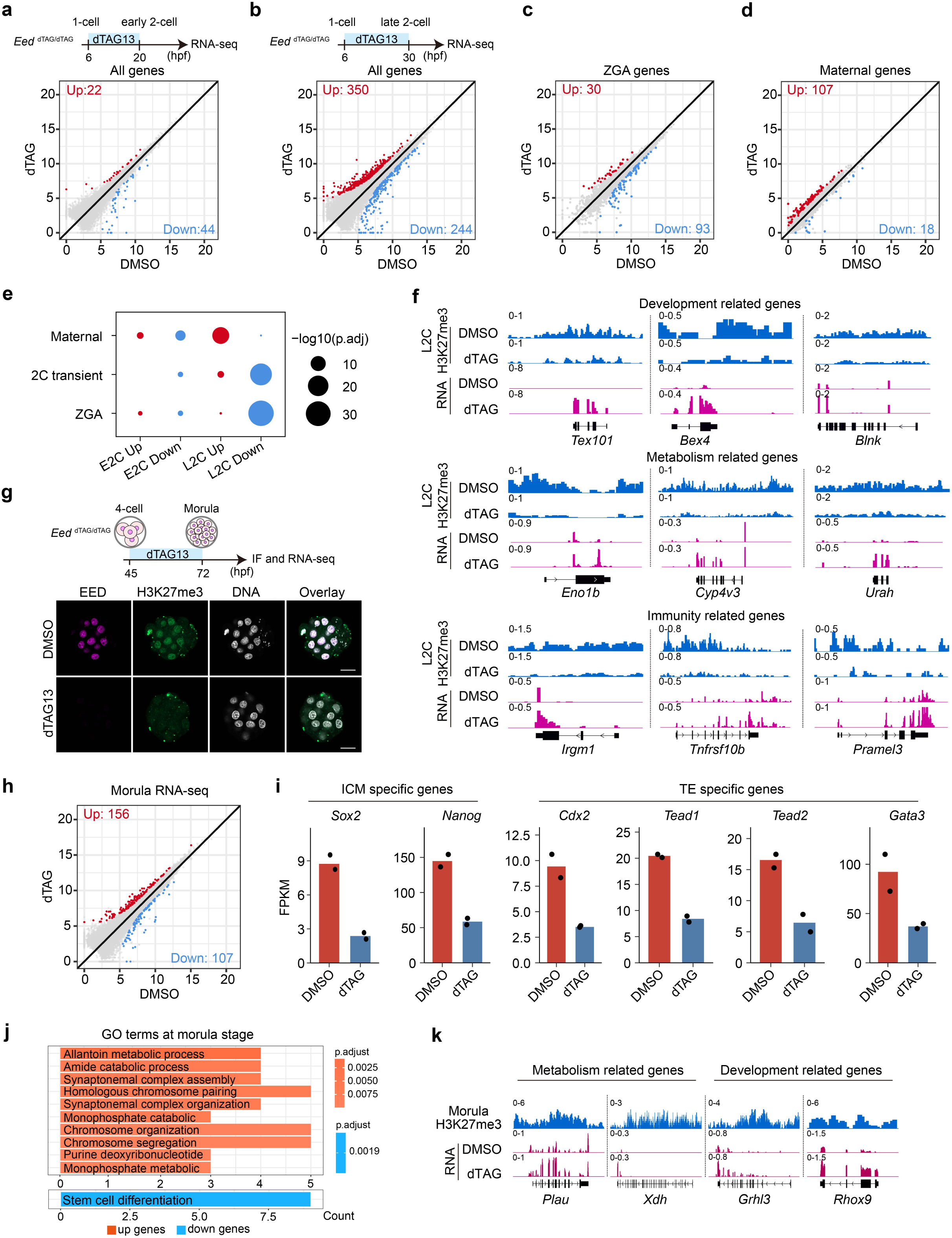
Effects of EED rapid depletion on maternal-to-zygotic transition and mid-preimplantation development. (**a-b**) Scatter plots comparing gene expression changes between DMSO and dTAG groups at early 2-cell (a) and late 2-cell (b) stages. Red and blue dots represent up- and down-regulated genes in the dTAG group, respectively. The cutoff used to define differentially expressed genes: fold change (FC) >= 2, adjusted *p*-value < 0.05, and FPKM >= 1. (**c-d**) Scatter plots comparing expression changes of the major ZGA genes (c) and the maternal genes (d) between DMSO and dTAG groups at late 2-cell stage. ZGA (n = 2,773) and maternal genes (n = 3,701) were defined. (**e**) Bubble plot showing the overlap between differentially expressed genes and the selected categories. *P*-values were calculated by one-sided hypergeometric test and adjusted by the Benjamini-Hochberg procedure. (**f**) Genome browser views of example genes that are enriched for H3K27me3 in late 2-cell embryos and are de-repressed upon rapid EED depletion. (**g**) Immunostaining images showing rapid depletion of EED and H3K27me3 in morulae after dTAG13 treatment starting from 4-cell stage. DNA, Hoechst 33342. Scale bar, 20 μm. Each experiment was independently repeated three times with similar results. One representative result is shown. (**h**) Scatter plot comparing gene expression changes between DMSO and dTAG groups at morula stage. (**i**) Bar plots showing the expression changes of representative ICM- and TE-specific genes in DMSO and dTAG groups at morula stage. Dots indicate individual RNA-seq biological replicates (n=2). (**j**) Gene ontology terms enriched for the differentially expressed genes in the dTAG group at morula stage. *P*-values were calculated by one-sided hypergeometric test and adjusted by the Benjamini-Hochberg procedure. (**k**) Genome browser views of example genes that are enriched for H3K27me3 in morulae and are de-repressed upon rapid EED depletion.

To identify potential direct targets of EED, the up-regulated genes were compared to the late 2-cell H3K27me3 with or without EED depletion (**Extended Data Fig. 2f, g**). We identified a few developmental genes (*e.g., Tex101* and *Bex4*), metabolic genes (*e.g., Eno1b* and *Cyp4v3*), and immune genes (*e.g., Irgm1* and *Tnfrsf10b*) that are enriched for H3K27me3 and de-repressed upon acute EED depletion (**Fig. 2f**). These data support that H3K27me3 is retained at some developmental genes and is essential for keeping them repressed during ZGA. Notably, the maternal gene *Ccnb3*, whose overexpression has been shown to cause ZGA failure ^39^, exhibits a mild upregulation upon PRC2 loss and is enriched for H3K27me3 (**Extended Data Fig. 2h, i**). These observations raise the possibility that *Ccnb3* may contribute to transcriptional dysregulation associated with the observed ZGA defects.

### PRC2 regulates lineage-specific genes during mid-preimplantation development

Since acute EED depletion affected both ICM and TE cell numbers in blastocysts (**Fig. 1i-k**), we next sought to understand how EED/PRC2 may regulate the first lineage specification. To avoid the potential secondary effects caused by EED loss during ZGA (**Fig. 2a**), *Eed^dTAG/dTAG^* embryos were cultured with dTAG13 from 4-cell (45 hpf) to morula stage (72hpf), when cells have undergone compaction (**Fig. 2g**). Immunostaining revealed that both EED and H3K27me3 were completely depleted at morula stage (**Fig. 2g**). Comparative transcriptome analyses identified 156 and 107 genes were up- and down-regulated, respectively, in response to dTAG13 treatment (**Fig. 2h, Extended Data Fig. 2j and Supplementary Table 2**). Interestingly, the well-known pluripotency genes (*i.e., Sox2* and *Nanog*) and trophoblast genes (*i.e., Cdx2* and *Gata3*) were downregulated in the dTAG group (**Fig. 2i**). Indeed, the down-regulated genes were enriched for gene ontology (GO) term “stem cell differentiation” (**Fig. 2j**). The up-regulated genes were enriched for “allantoin metabolic process”, which should normally occur in later development (**Fig. 2j, k**). In addition, a subset of totipotent genes became reactivated at the morula stage upon EED degradation (**Extended Data Fig. 2k**). Taken together, these data support that PRC2-mediated transcriptional silencing during mid-preimplantation development is required to ensure proper lineage-specific gene expression.

### PRC2 regulates EPI formation by preventing premature activation of developmental genes

Given that EED depletion impairs blastocyst formation (**Fig. 1g-h**) and regulates lineage-specific genes during compaction (**Fig. 2i**), we asked whether PRC2 could also regulate the second lineage specification. To avoid indirect effects caused by EED loss in early- and mid- preimplantation development, *Eed^dTAG/dTAG^*embryos were cultured with dTAG13 starting from morula (72 hpf) or mid-blastocyst (96 hpf) to late-blastocyst stage (112 hpf) (**Fig. 3a**). Both treatment windows nearly completely depleted both EED and H3K27me3 (**Fig. 3b**). Remarkably, acute EED degradation causes decrease in the percentage of EPI but not PrE cells (**Fig. 3c, d**), supporting that EED plays an important role in regulating EPI formation.

**Figure 3.**
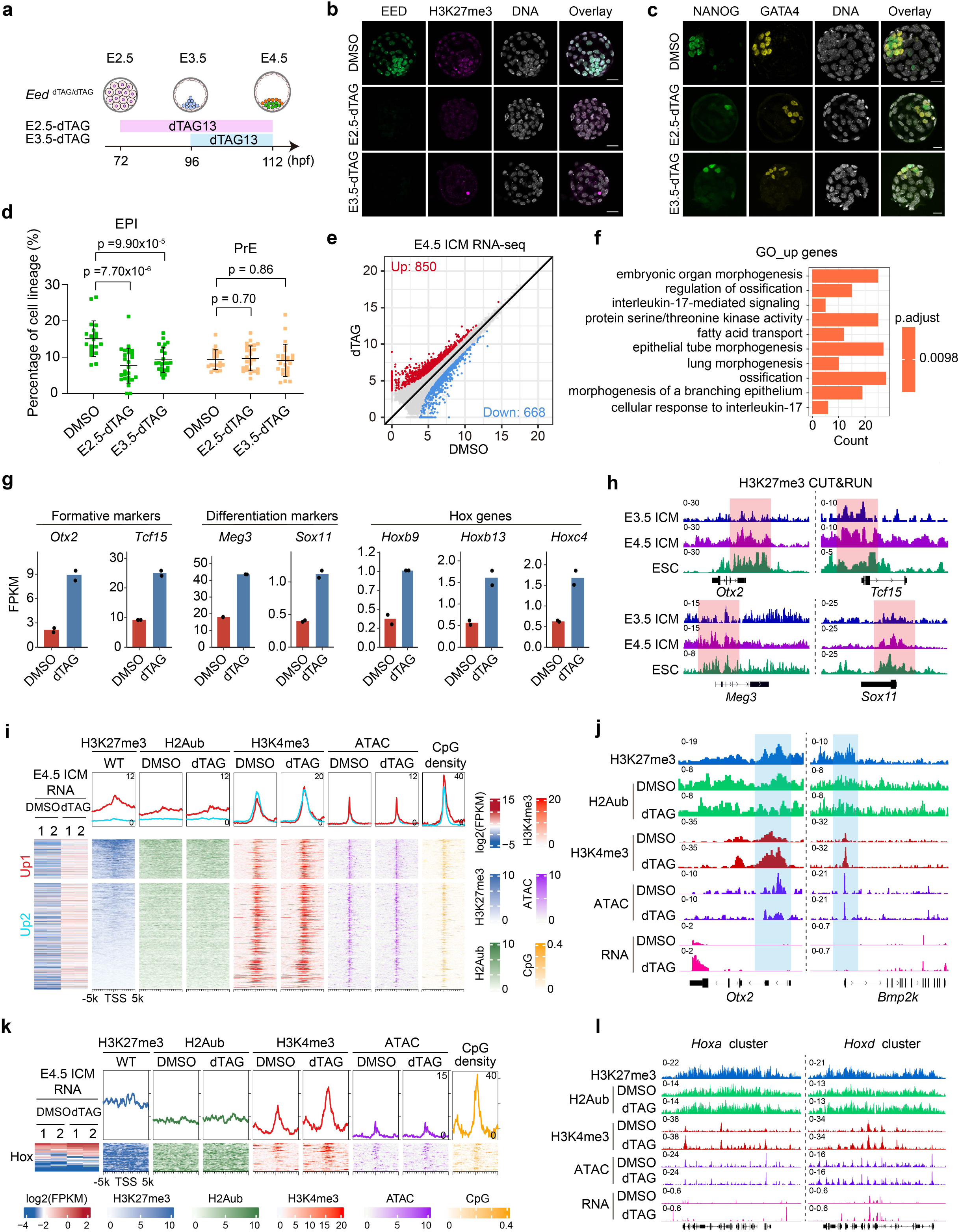
Effects of EED rapid depletion on lineage specification in late blastocysts. (**a**) Schematic showing the dTAG13 treatment strategies for evaluating EED function in lineage specification in late blastocysts. (**b**) Immunostaining images showing the rapid depletion of EED and H3K27me3 in late blastocysts after dTAG13 treatment starting from morula (E2.5-dTAG) or early blastocyst (E3.5-dTAG) stages. DNA, Hoechst 33342. Scale bar, 20 μm. (**c**) Immunostaining images showing the EPI (NANOG^+^/GATA4^-^) and PrE (GATA4^+^/NANOG^-^) cells at late blastocyst stage in DMSO and dTAG groups. EPI: epiblast; PrE: primitive endoderm. Hoechst 33342. Scale bar, 20 μm. (**d**) Quantifications of EPI and PrE cells of panel c (DMSO, n=20; E2.5-dTAG, n=26; E3.5-dTAG, n=24). n represent the embryo numbers. *p* values were calculated with Student’s *t*-test (two-sided). The data are presented as mean values ± SD. Experiments were repeated three times. (**e**) Scatter plots comparing gene expression changes between DMSO and dTAG groups for E4.5 ICM cells. Red and blue dots represent up-and down-regulated genes in the dTAG group, respectively. (**f**) Gene ontology terms enriched for the differentially expressed genes in the dTAG group for E4.5 ICM cells. *p*-values were calculated by one-sided hypergeometric test and adjusted by the Benjamini-Hochberg procedure. (**g**) Bar plots showing the expression changes of representative formative markers, differentiation markers, and *Hox* genes in DMSO and dTAG groups for E4.5 ICM cells. Dots indicate individual RNA-seq biological replicates (n=2). (**h**) Genome browser views of example genes that are enriched with H3K27me3 in E3.5/E4.5 ICM and are de-repressed upon rapid EED depletion. (**i**) Meta plots and heatmaps showing the changes of RNA, H2Aub, H3K4me3, and ATAC signals in E4.5 ICM upon EED depletion. The up-regulated genes in the dTAG group are classified into two groups based on whether H3K27me3 is enriched. (**j**) Genome browser views showing the changes of H2Aub, H3K4me3, ATAC, and RNA signals at *Otx2* and *Bmp2k* loci in E4.5 ICM cells upon rapid EED depletion. (**k**) Meta plots and heatmaps showing the changes of RNA, H2Aub, H3K4me3, and ATAC signals at *Hox* loci in E4.5 ICM upon EED depletion. (**l**) Genome browser views showing the changes of H2Aub, H3K4me3, ATAC, and RNA signals at the *Hoxa* and *Hoxc* loci in E4.5 ICM cells upon rapid EED depletion. Source numerical data are available in source data.

To understand the underlying mechanisms, E4.5 ICM cells were isolated by immuno-surgery ^40^ and subjected to RNA-seq analyses. Comparative analyses revealed 850 and 668 up- and down-regulated genes with EED depletion, respectively (**Fig. 3e, Extended Data Fig. 3a and Supplementary Table 3**). GO term analyses revealed that the up-regulated genes are enriched for developmental processes such as “embryonic organ morphogenesis” (**Fig. 3f**). While EED depletion at this stage does not affect pluripotency gene markers such as *Nanog* and *Sox2* (**Extended Data Fig. 3b**), but up-regulate the formative pluripotency marker genes, such as *Otx2* and *Tcf15* (**Fig. 3g**). This data suggests that PRC2 prevents premature activation of genes responsible for naïve- to-formative pluripotency transition. Premature activation of *Otx2* and *Tcf15* may contribute to the defects in EPI formation ^41, 42^. In addition, differentiation markers (*i.e., Meg3* and *Sox11*) and Hox genes, which supposed to be expressed at primed EPI or gastrulation stages were up-regulated when EED was depleted (**Fig. 3g**). Importantly, all these genes are enriched for H3K27me3 in control but lost upon EED depletion, suggesting that they are directly repressed by H3K27me3 (**Fig. 3h and Extended Data Fig. 3c, d**). Collectively, these data indicate that PRC2 is essential for EPI formation by preventing premature activation of developmental genes.

### PRC2 constrains H3K4me3 levels in E4.5 ICM

We next attempted to connect chromatin changes and the transcriptome dysregulations caused by EED depletion in E4.5 ICM. Analyses of the H2Aub indicated that EED depletion did not affect global H2Aub (**Fig. 3i, j and Extended Data Fig. 3e, f**). This observation is consistent with our previous observation that non-canonical PRC1 deposits the majority of H2Aub independent of PRC2 from oocytes to 4-cell embryos^43^. Notably, EED depletion caused an increase in H3K4me3 at PRC2 targeted genes, where H3K27me3 was highly enriched (“Up1” group), but to a much less extent for the H3K27me3 lowly enriched genes (“Up2” group) (**Fig. 3i, j and Extended Data Fig. 3f**), suggesting a direct effect on H3K4me3. However, loss of EED did not affect chromatin accessibility as revealed by ATAC-seq (**Fig. 3i and Extended Data Fig. 3f)**, suggesting that the increase in H3K4me3 was independent of chromatin accessibility. In addition, *Hox* genes also showed an increase in H3K4me3 and RNA levels in the dTAG group (**Fig. 3k, l and Extended Data Fig. 3f**). These data suggest that PRC2 constrains H3K4me3 and prevents activation of PRC2 targets in E4.5 ICM.

To determine whether transient loss of PRC2 can exert lasting effects on chromatin state, we depleted EED from zygotes to 4-cell embryos by treating embryos with dTAG13 during this time window, then washing out for further development. E4.5 ICMs were collected for CUT&RUN analysis (**Extended data Fig. 3g**). We found that H3K4me3 was modestly increased with a partial restoration of H3K27me3, suggesting that H3K4me3 recovery occurs in parallel with changes in H3K27me3 (**Extended Data Fig. 3h**). Given that H3K27me3 was not fully restored at E4.5, it remained unclear whether the increase in H3K4me3 in E4.5 ICM resulted from transient H3K27me3 loss at the 4-cell stage or reflected stage-specific regulation at E4.5. To distinguish between these two possibilities, we examined loci marked by H3K27me3 specifically at the 4-cell stage. We found that transient depletion of H3K27me3 at the 4-cell stage had only a minor effect on global H3K4me3 levels at E4.5 (**Extended Data Fig. 3i**). However, genes marked by H3K27me3 in E4.5 ICM exhibited an increase in H3K4me3 level, consistent with partial H3K27me3 loss contributing to H3K4me3 increase (**Extended Data Fig. 3i**).

### PRC2 plays critical role for bivalency establishment in E4.5 ICM

The fact that both H3K27me3 and H3K4me3 are enriched in developmental genes like *Otx2* and *Bmp2k* at E4.5 ICM (**Fig. 3i, j**) suggests a bivalency state ^44^. Previous studies suggest that bivalency is largely absent or infrequent from developmental genes in preimplantation embryos (up to E3.5 ICM) but appears after implantation (E5.5-E6.5 EPI) ^34, 45^. This inconsistency prompted us to further analyze the bivalent genes throughout pre- and post-implantation development (Methods). This analysis confirmed the largely absence of bivalency before E4.5 ICM, with only 272 and 364 bivalent promoters in 8-cell and E3.5 ICM, respectively (**Fig. 4a-c**). However, a major transition occurs between E3.5 to E4.5 with a ∼4-fold increase (1,457 vs. 364) of bivalent genes by E4.5 ICM (**Fig. 4c and Supplementary Table 4**). The number of bivalent genes further increased to 4,382 by E6.5 EPI. Consistent with the notion that bivalency is to poise for activation, the bivalent genes identified are mostly not expressed or lowly expressed in E4.5 ICM and E6.5 EPI (**Extended Data Fig. 4a, b and Supplementary Table 4**).

**Figure 4.**
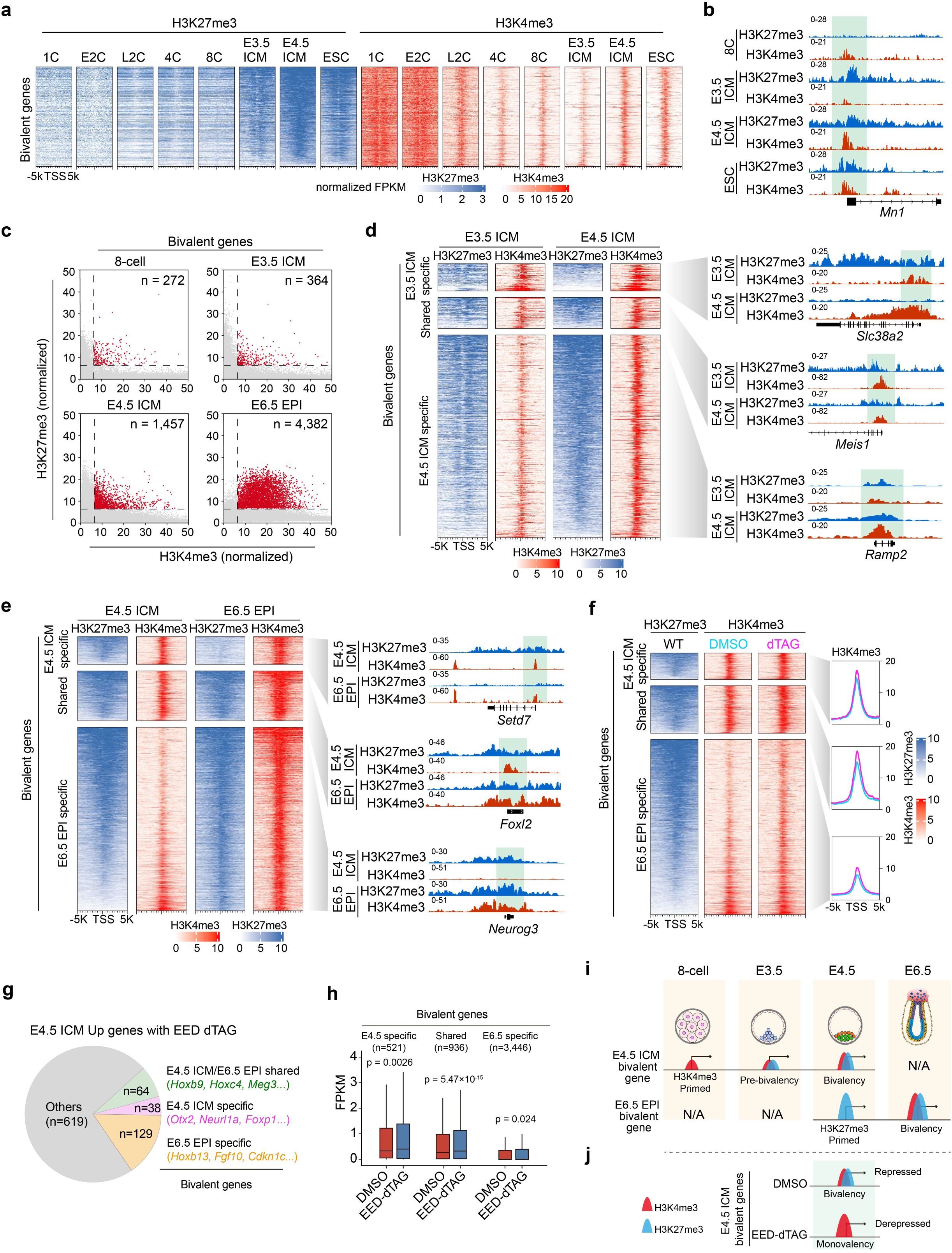
Effects of EED rapid depletion on bivalency formation in pre- and early post-implantation embryos. (**a**) Dynamics of H3K27me3 and H3K4me3 at bivalent genes identified in E4.5 ICM (n = 1457, Methods) throughout preimplantation development. (**b**) Genome browser views of H3K27me3 and H3K4me3 dynamics at an example bivalent locus from 8-cell to E4.5 ICM and ESCs. (**c**) Scatter plot showing the number of bivalent genes (red dots) at the indicated developmental stages. (**d**) Dynamics of H3K27me3 and H3K4me3 at bivalent loci during the E3.5 ICM to E4.5 ICM transition. Left panel showing heatmaps of these two histone modifications in E3.5 ICM and E4.5 ICM cells. All bivalent loci are classified into three groups: “E3.5 ICM specific”, “Shared”, and “E4.5 ICM specific”. Right panel showing genome browser views of example genes for each group. (**e**) Dynamics of H3K27me3 and H3K4me3 at bivalent loci during the E4.5 ICM to E6.5 EPI transition. Left panel showing heatmaps of these two histone modifications in E4.5 ICM and E6.5 ICM cells. All bivalent loci are classified into three groups: “E4.5 ICM specific”, “Shared”, and “E6.5 ICM specific”. Right panel showing genome browser views of example genes for each group. (**f**) Effects of EED rapid depletion on H3K4me3 at bivalent loci in E4.5 ICM. Left panel showing heatmaps of H3K27me3 and the changes of H3K4me3 upon EED depletion. Right panel showing the metaplots that summarizing average H3K4me3 signals at the bivalent genes. (**g**) Pie chart showing the bivalent genes that are up regulated in E4.5 ICM upon EED depletion. (**h**) Boxplot showing the RNA expression changes of different groups of bivalent genes upon EED depletion in E4.5 ICM cells. *P*-values are calculated with two-sided Wilcoxon signed rank test. In the boxplot, the central band represents the median. The lower and upper edges of the box represent the first and third quartiles, respectively. The whiskers of the boxplot extend to 1.5 times interquartile range (IQR). (**i**) Schematic model summarizing the dynamics of H3K4me3 and H3K27me3 at bivalent loci during the 8-cell to E3.5/E4.5 ICM and E4.5 ICM to E6.5 EPI transitions. (**j**) Schematic model summarizing the impact of EED rapid depletion on H3K4me3 and H3K27me3 at bivalent genes in E4.5 ICM cells.

We next sought to understand how these major transitions of bivalency took place in E4.5 ICM and E6.5 EPI. Clustering analyses revealed that gain of bivalency in E4.5 ICM is mainly due to the further increase in both H3K27me3 and H3K4me3 (**Fig. 4d**, “E4.5 ICM specific”). Note that this group of genes already showed moderate and/or weak enrichment of both modifications in E3.5 ICM. In contrast, only a small percentage (194 out of 1457, 13.3%) of bivalent genes are inherited or maintained from E3.5 to E4.5 ICM (**Fig. 4d**, “Shared”). Unlike the E3.5 to E4.5 ICM transition, the gain of bivalency from E4.5 ICM to E6.5 EPI was mainly due to the increase of H3K4me3 and the inheritance of H3K27me3 (**Fig. 4e**). The bivalent genes of all the groups are enriched for similar GO terms such as cell fate commitment and embryonic organ morphogenesis (**Extended Data Fig. 4c, d**). Consistent with earlier analyses (**Fig. 3i**), EED depletion caused increased H3K4me3 at bivalent genes in E4.5 ICM, even including those that normally gain H3K4me3 in E6.5 EPI (**Fig. 4f**, “E6.5 EPI specific”). Furthermore, the increase of H3K4me3 at bivalent genes correlated with their increased expression (**Fig. 4g**), with a stronger effect for genes already show high levels of H3K4me3 in E4.5 ICM (“E4.5-specific” and “Shared”) (**Fig. 4h**).

Collectively, these analyses revealed the different mechanisms underlying bivalency acquisition in ICM and EPI (**Fig. 4i)**. These data also demonstrate that PRC2 constrains H3K4me3 and transcription levels at bivalent genes in E4.5 ICM (**Fig. 4j**).

### PRC2 restricts PGC numbers by repressing BMP4 signaling through a lineage crosstalk

We further explored the role of PRC2 in regulating bivalency genes in post-implantation embryos. To this end, we first confirmed that the dTAG system can rapidly and efficiently degrade EED (∼4 hr) in *ex vivo* cultured post-implantation embryos (**Extended Data Fig. 5a, b**) and then tested the degradation efficiency *in vivo*. To minimize the impact of EED depletion on the pregnant mice, *Eed^dTAG/+^* female mice were used to mate with *Eed^dTAG/dTAG^* male mice, and subjected to intraperitoneal (IP) injection of dTAG^V^-1. The embryos were collected for immunostaining at E6.5 (**Fig. 5a**) with the *Eed^dTAG/+^* embryos in the same litter served as a control. Immunostaining analyses indicated that both EED and H3K27me3 were near completely depleted after ∼12-hr dTAG^V^-1 treatment (**Fig. 5a**). Similarly, administration of dTAG^V^-1 at E6.5 also led to near complete degradation of both EED and H3K27me3 by E7.5 (**Fig. 5a**). Thus, the Eed-dTAG system allows efficient and inducible degradation of PRC2 in early post-implantation embryos *in vivo*.

**Figure 5.**
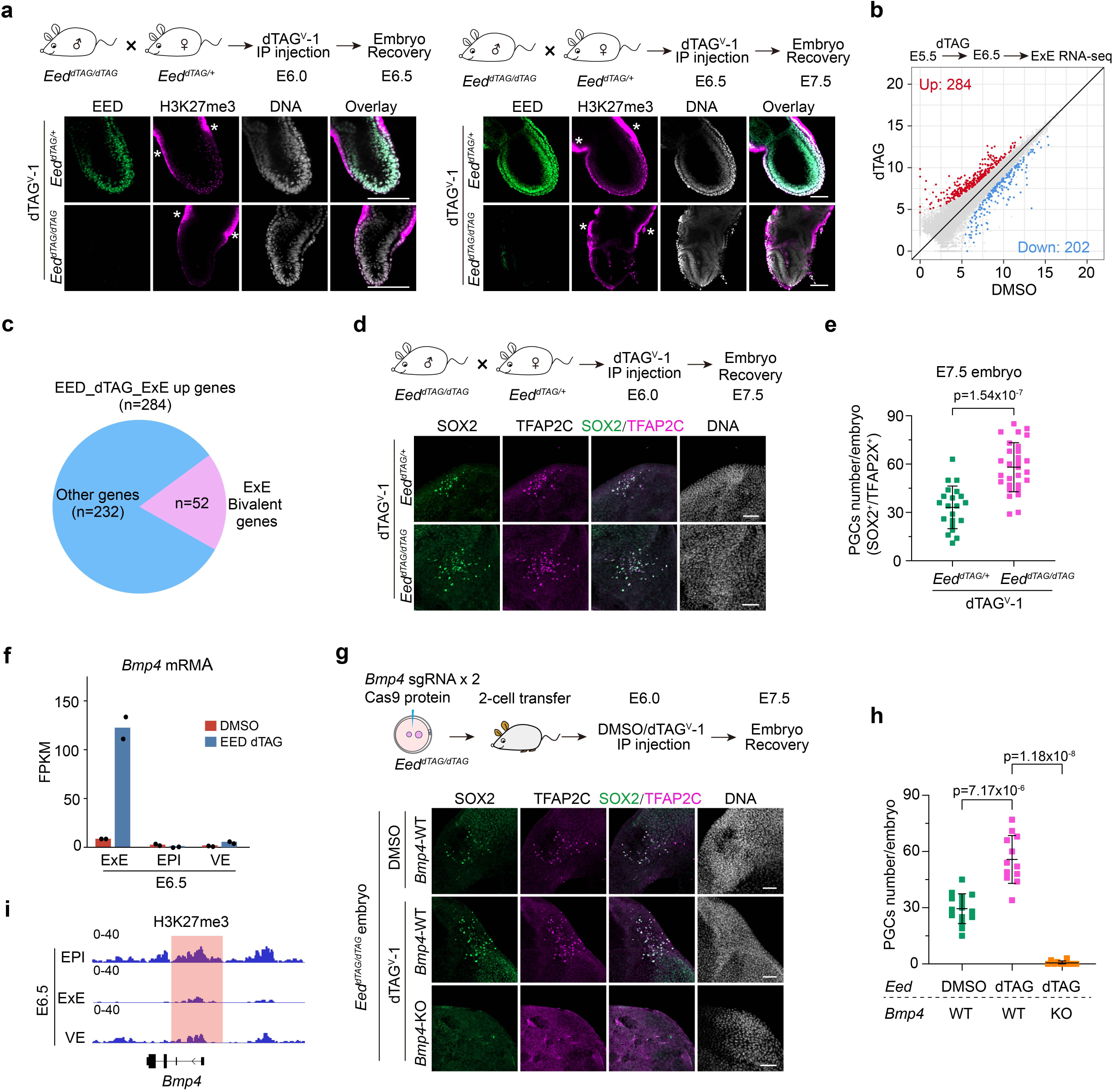
EED depletion during gastrulation causes primordial germ cells (PGCs) increase through BMP4 signaling. (**a**) Immunostaining images showing the rapid depletion of EED and H3K27me3 in post-implantation embryos (*i.e.,* E6.5 or E7.5) following administration of dTAG^V^-1 by Intraperitoneal (IP) injection. Hoechst 33342. Scale bar, 100 μm. *showed the non-specific staining of H3K27me3. Each experiment was independently repeated at least three times with similar results. One representative result is shown. **(b)** Scatter plot comparing gene expression changes between DMSO and dTAG groups in E6.5 ExE. *Eed^dTAG/+^* female mice were used to mate with *Eed^dTAG/dTAG^* male mice, and subjected to IP injection of DMSO or dTAG^V^-1 at E5.5. The embryos were dissected for RNA-seq. **(c)** Pie chart showing the number of ExE bivalent genes in the up-regulated genes upon EED depletion. (**d**) Immunostaining images showing the increased PGCs (SOX2^+^/TFAP2C^+^) in E7.5 embryos after EED depletion. Hoechst 33342. Scale bar, 100 μm. (**e**) Quantification of PGCs in panel d (*Eed^dTAG/+^*, n=21; *Eed^dTAG/dTAG^*, n=29). *p* values were calculated with Student’s *t*-test (two-sided). The data are presented as mean values ± SD. Experiments were repeated three times. (**f**) Bar plot showing the RNA expression changes of *Bmp4* after EED depletion in E6.5 EPI, ExE, and VE. Dots indicate individual RNA-seq biological replicates (n=2). (**g**) Top panel: schematic showing the experimental design to evaluate how acute *Bmp4* KO may affect PGCs with or without dTAG-mediated EED depletion in E7.5 embryos; Bottom panel: immunostaining images showing the number of PGCs in E7.5 embryos following EED single depletion or EED/BMP4 double depletion. Hoechst 33342. Scale bar, 100 μm. (**h**) Quantification of PGCs in panel g (*Eed*-DMSO/*Bmp4*-WT, n=15; *Eed*-dTAG/*Bmp4*-WT, n=12; *Eed*-dTAG/*Bmp4*-KO, n=11). *p* values were calculated with Student’s *t*-test (two-sided). The data are presented as mean values ± SD. Experiments were repeated three times. (**i**) Genome browser view of H3K27me3 at the *Bmp4* locus in E6.5 EPI, ExE and VE lineages. Source numerical data are available in source data.

For the RNA-seq, *Eed^dTAG/+^* female mice were used to mate with *Eed^dTAG/dTAG^* male mice, and subjected to intraperitoneal (IP) injection of DMSO or dTAG^V^-1 at E5.5. We then dissected E6.5 EPI, ExE and visceral endoderm (VE) from DMSO or dTAG-treated embryos for RNA-seq analyses (**Extended Data Fig. 5c**).

Unexpectedly, acute EED depletion had limited impact on EPI and VE transcriptomes (**Extended Data Fig. 5d and Supplementary Table 5**). However, more genes, including bivalent genes (52 out of 284) are up-regulated in ExE lineage after EED degradation (**Fig. 5b, c, Extended Data Fig. 5e and Supplementary Table 5**). Further analysis showed that the global bivalent gene expression is increased with EED degradation (**Extended Data Fig. 5f and Supplementary Table 4**). The GO terms enriched in the up-regulated genes in EED-depleted E6.5 ExE include developmental processes such as “ossification” and “mesenchyme development” (**Extended Data Fig. 5g**), suggesting PRC2 also repress development genes in ExE lineage. These results indicate that PRC2 is necessary for silencing bivalent genes and development genes in the ExE lineage.

Given that the ExE lineage derived BMP4 is essential for PGC induction in a dosage-dependent manner ^46–48^, and that loss of PRC2 lead to overproduction of PGCs (SOX2^+^/TFAP2C^+^) (**Fig. 5d, e**) ^37^, we hypothesized that PRC2-mediated repression of *Bmp4* in the ExE lineage might regulate PGCs specification. Indeed, *Bmp4* mRNA level increased by ∼15 fold upon EED depletion in E6.5 ExE, but not in EPI or VE (**Fig. 5f**). To determine whether the increased *Bmp4* level in the ExE is responsible for the expanded PGC fate in EED-depleted embryos (**Fig. 5d, e**), we knocked out *Bmp4* in *Eed^dTAG/dTAG^* embryos by zygotic CRISPR injection and assessed PGCs numbers at E7.5. Remarkably, *Bmp4* KO caused complete loss of PGCs even in EED*-*depleted embryos (**Fig. 5g, h**). Given that PGCs are derived from the EPI lineage, and the *Bmp4* is upregulated in the ExE lineages in response to EED degradation, these data support that EED restricts PGC numbers in EPI through a lineage crosstalk between ExE and EPI. Interestingly, CUT&RUN analyses showed negligible H3K27me3 enrichment at the *Bmp4* locus (**Fig. 5i and Extended Data Fig. 5h**) indicating that PRC2 represses *Bmp4* in the ExE lineage likely through an indirect mechanism.

### PRC2 represses *Bmp4* through repressing the bivalent gene *Esrrb* in ExE

To investigate how PRC2 activity in ExE may regulate PGCs specification in EPI, we identified a list of transcription factors that are enriched for both H3K27me3 and H3K4me3 marks in E6.5 ExE and are up-regulated upon EED depletion in the same lineage (**Fig. 6a**). Of these candidates, *Esrrb* has been shown to induce *Bmp4* expression in E6.5 ExE and is critical for PGC specification^49^. Interestingly, *Esrrb* is a bivalent gene enriched for both H3K4me3 and H3K27me3 in the ExE lineage (**Fig. 6b and Extended Data Fig. 5e**).

**Figure 6.**
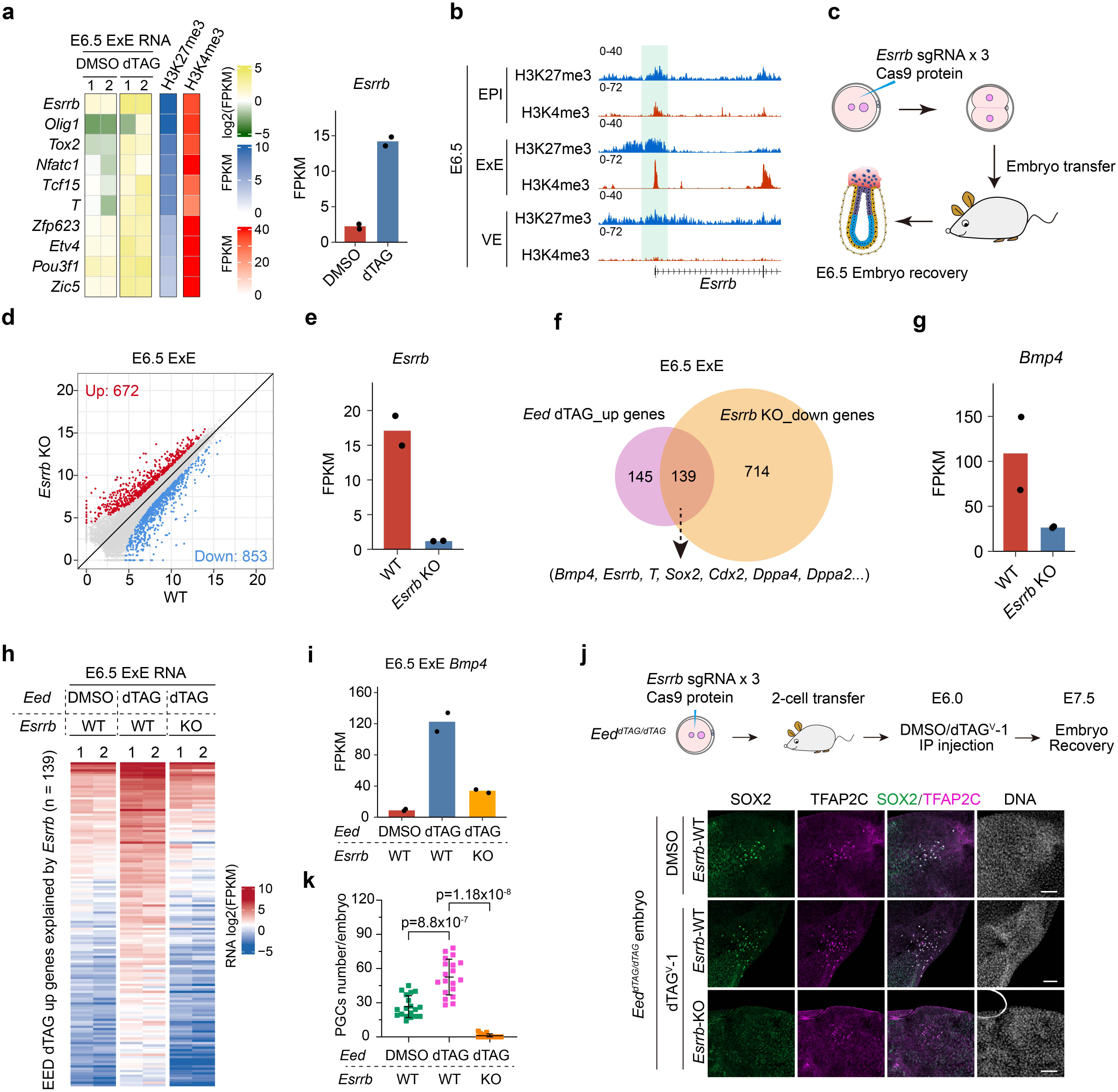
EED suppresses PGC fate by directly repressing *Esrrb*. (**a**) Heatmap showing RNA, H3K27me3 and H3K4me3 levels of the candidate bivalent TFs that may be involved in PGC specification in E6.5 ExE. Right panel showing bar plot of *Esrrb* expression without and with EED degradation. Dots indicate individual RNA-seq biological replicates (n=2). (**b**) Genome browser views of H3K4me3 and H3K27me3 at the *Esrrb* locus in E6.5 EPI, ExE, and VE lineages. (**c**) Schematic showing the experimental design for acute *Esrrb* KO by zygotic CRISPR injection. (**d**) Scatter plot comparing gene expression changes between *Esrrb* wild type (WT) and KO E6.5 ExE. (**e**) Bar plot showing the RNA expression changes of *Esrrb* in *Esrrb* KO E6.5 ExE. The adjusted *p*-value was calculated by DESeq2. Dots indicate individual RNA-seq biological replicates (n=2). (**f**) Venn diagram showing the overlap of genes that are de-repressed upon EED depletion and genes that are down-regulated by *Esrrb* KO in E6.5 ExE. (**g**) Bar plot showing the RNA expression changes of *Bmp4* in E6.5 ExE after *Esrrb* KO. The adjusted *p*-value was calculated by DESeq2. Dots indicate individual RNA-seq biological replicates (n=2). (**h**) Heatmap illustrating the RNA expression changes of the 139 genes that are de-repressed by EED depletion but are down-regulated in E6.5 ExE of *Esrrb* KO. (**i**) Bar plot showing the RNA expression changes of *Bmp4* in E6.5 ExE following EED single depletion or EED/ESRRB double depletion. Dots indicate individual RNA-seq biological replicates (n=2). (**j**) Top panel: schematic showing the experimental design to evaluate how acute KO of *Esrrb* may affect PGC numbers with or without EED depletion in E7.5 embryos; Bottom panel: immunostaining images showing the number of PGCs in E7.5 embryos following EED single depletion or EED/ESRRB double depletion. Hoechst 33342. Scale bar, 100 μm. (**k**) Quantification of PGC numbers in panel j (*Eed*-DMSO/*Esrrb*-WT, n=19; *Eed*-dTAG/*Esrrb*-WT, n=19; *Eed*-dTAG/*Esrrb*-KO, n=14). *P* values were calculated with Student’s *t*-test (two-sided). The data are presented as mean values ± SD. Experiments were repeated three times. Source numerical data are available in source data.

To identify downstream targets of *Esrrb*, we acutely knocked out *Esrrb* by zygotic CRISPR injection and performed RNA-seq analyses in E6.5 ExE (**Fig. 6c**). Despite that depletion of *Esrrb* did not affect overall embryo morphology at this stage (**Extended Data Fig. 6a**), which is consistent with a previous study^49^, transcriptome analyses revealed 672 and 853 genes were respectively up- and down-regulated in *Esrrb* KO E6.5 ExE (**Fig. 6d, Extended Data Fig. 6b and Supplementary Table 6**). Down-regulation of *Esrrb* confirmed the successful KO of *Esrrb* in the embryos analyzed (**Fig. 6e**). The up-regulated genes are enriched for GO terms including “actin filament organization”, whereas the down-regulated genes are enriched for GO terms such as “formation of primary germ layer” (**Extended Data Fig. 6c, d**). Interestingly, 139 of the down-regulated genes in *Esrrb* KO were also up-regulated in response to EED depletion (**Fig. 6f and Extended Data Fig. 6e**), indicating that the up-regulated genes, such as *Bmp4,* in the EED-depleted embryos could be mediated by the de-repression of *Esrrb* (**Fig. 6a, f, g**).

To test this hypothesis, we acutely depleted *Esrrb* in *Eed^dTAG/dTAG^*embryos by zygotic CRISPR injection followed by RNA-seq analyses in E6.5 ExE samples. As expected, upregulation of the 139 genes in the EED-depleted embryos were near completely reversed by *Esrrb* KO (**Fig. 6h**). Notably, the increased *Bmp4* RNA level upon EED depletion can be largely rescued by *Esrrb* KO (**Fig. 6i**). Nonetheless, we next evaluated to what extent *Esrrb* KO may reverse the expanded PGC phenotype in the EED-depleted embryos (**Fig. 5d, e**). Like *Bmp4* KO (**Fig. 5g, h**), *Esrrb* KO also greatly reduced the PGC numbers in EED-depleted E7.5 embryos (**Fig. 6j, k**). Collectively, these data indicate that PRC2 suppresses *Bmp4* and PGC fate largely through direct repressing *Esrrb* in the ExE lineage, highlight the lineage crosstalk-dependent functions of PRC2 in regulating PGCs formation.

### *Esrrb* regulates its downstream targets primarily by binding to their enhancers

Having demonstrated that *Esrrb* is a direct target of PRC2, and ESRRB promotes PGC fate by activating *Bmp4* signaling (**Fig. 6**), we next sought to understand how ESRRB regulates its downstream target genes at the chromatin level. To this end, we first performed ESRRB CUT&RUN analyses in E6.5 and E7.0 ExE lineage (**Fig. 7a and Extended Data Fig. 7a**). ESRRB signals are markedly reduced in the knockout samples, confirming the specificity of the signals (**Fig. 7a)**. In addition, ESRRB co-binding with its co-factor SOX2 (**Extended Data Fig. 7b, c**)^50^, further confirm the reliability of ESRRB signals. Detailed analyses revealed that the ESRRB peaks are mostly located at introns and distal intergenic regions, with only a small percentage of peaks found at promoter regions (**Extended Data Fig. 7d**). This observation indicates that ESRRB mainly bind to putative enhancer regions. Indeed, ESRRB bound to well characterized enhancers of *Bmp4*, *Cdx2,* and *Sox2* (**Fig. 7b**). Importantly, all these genes are downregulated upon *Esrrb* KO in E6.5 ExE samples (**Fig. 7b)**, supporting that ESRRB promotes expression of these genes by activating their enhancers.

**Figure 7.**
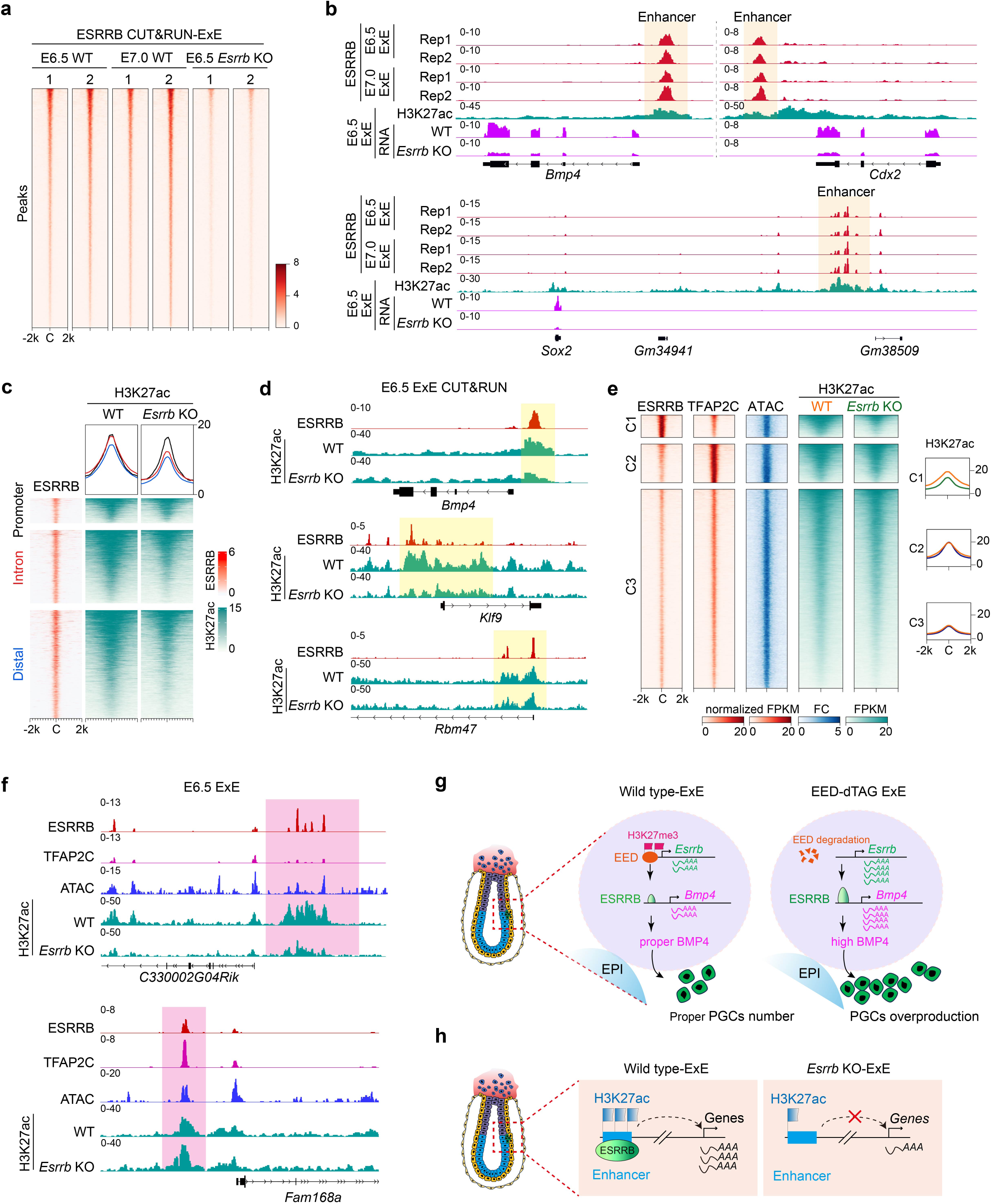
ESRRB regulates gene expression in ExE through binding to enhancers. (**a**) Heatmap illustrating ESRRB CUT&RUN signal enrichment across wild-type (WT) E6.5 and E7.0 ExE, as well as the *Esrrb*- KO E6.5 ExE lineage. Two biological replicates for each stage are shown. (**b**) Genome browser views of ESRRB binding and RNA signals at the indicated genomic loci. H3K27ac marks active enhancer location. (**c**) Heatmap showing the H3K27ac changes caused by *Esrrb* KO at ESRRB-bound promoters, introns, and distal regions in E6.5 ExE. (**d**) Genome browser views of ESRRB binding and H3K27ac changes at the indicated genomic loci. (**e**) Left panel: heatmaps showing ATAC signal and the H3K27ac changes caused by *Esrrb* KO at ESRRB and TFAP2C co-bound regions. The peaks were classified into three clusters based on ESRRB and TFAP2C enrichment levels. (**f**) Genome browser views of ESRRB and TFAP2C binding, ATAC signal, and H3K27ac level at the indicated genomic loci. (**g**) Diagram summarizing the EED-ESRRB-BMP4 axis in regulating PGC specification during gastrulation. (**h**) Diagram illustrating that ESRRB regulates its target genes by activating enhancers.

Since one hallmark of enhancer is the enrichment of H3K27ac ^51^, we next investigated how loss of ESRRB may affect H3K27ac. To this end, we performed H3K27ac CUT&RUN analyses in E6.5 ExE and found that H3K27ac levels at ESRRB-bound distal/intron regions (i.e., *Bmp4*, *Klf9*, *Wnt6*, *Sox21*, *Fbxo21*) are greatly reduced upon *Esrrb* KO (**Fig. 7c, d and Extended Data Fig. 7e**). This observation confirms a role of ESRRB in establishing enhancer function in regulating formative transition and trophoblast stem cells ^52–54^. In contrast, the effect of *Esrrb* KO on H3K27ac levels at ESRRB-bound promoters is much milder (i.e., *Rbm47*, **Fig. 7c, d**).

To understand how ESRRB may synergistically activate enhancers with other transcription factors in ExE, we next focused on TFAP2C, a critical TF for ExE development ^55, 56^ with many shared targets with ESRRB (**Fig. 7e**). Interestingly, clustering analyses revealed that a subset of the co-bound peaks exhibited biased enrichment of these two transcription factors (i.e., cluster 1 and cluster 2 in **Fig. 7e**), indicating that these two TFs may have differential preference to different enhancer elements. Interestingly, ESRRB only regulates H3K27ac levels at peaks where it had a stronger enrichment than TFAP2C (**Fig. 7e, f**). It’s possible that TFAP2C may compensate the enhancer activity upon ESRRB KO at the peaks with a biased enrichment of TFAP2C. Thus, these data suggest that ESRRB regulates H3K27ac deposition mainly at its target gene enhancers, but not promoters, in E6.5 ExE. Additionally, at the ESRRB predominate enhancers, ESRRB is critical for gene activation independent of TFAP2C.

Collectively, these data support our notion that PRC2 plays a critical role in restricting PGCs number through a PRC2-ESRRB-BMP4 regulatory axis (**Fig. 7g**), and ESRRB regulates gene expression mainly by modulating enhancer activity (**Fig. 7h**).

## Discussion

In this study, we systematically evaluated the function of PRC2 in mouse early embryogenesis by using a EED-dTAG mouse model. Such targeted protein degradation system has two major advantages compared to conventional zygotic and maternal KOs. First, it enables depletion of maternal EED without using oocyte-specific Cre lines (*i.e., Gdf9-Cre* and *Zp3-Cre*). The germline conditional KO lines activate *Cre* expression at early stage of oogenesis (*i.e.,* postnatal day 3-5)^57^ and may cause accumulative compensatory and/or secondary effects in growing oocytes. Indeed, loss of PRC2 results in de-repression of certain Polycomb targets in oocytes, although the phenotypes are much milder than those of the PRC1 conditional KO ^24, 58, 59^. Second, the dTAG system allows for rapid depletion of EED at different developmental stages to reveal stage-specific PRC2 functions, which is not feasible when using conventional KO approaches. Using this strategy, we are able to uncover previously underappreciated stage-specific roles of PRC2 in regulating MZT, cell lineage-specification, and bivalency acquisition during pre- and peri-implantation development. We further elucidated the mechanism underlying how PRC2 controls proper PGC numbers through a PRC2-ESRRB-BMP4 regulatory axis and a lineage crosstalk.

Previous zygotic and maternal KOs have demonstrated important functions of PRC2 in early embryogenesis. The key conclusions from these studies are: 1) zygotic PRC2 is required for proper gastrulation, and 2) maternal PRC2 is critical for post-implantation development. Specifically, maternal PRC2 KO using *Gdf9-Cre* or *Zp3-Cre* causes defective imprinted X-chromosome inactivation (XCI), loss of non-canonical imprinting, partial lethality after implantation, and placental hyperplasia ^26, 27, 29–31, 40, 60^. In contrast to the earlier studies emphasizing the roles of PRC2 in post-implantation development, our study revealed a previously not appreciated PRC2 function in regulating pre-implantation development. We showed that depletion of EED results in abnormal gene expression as exemplified by the reduced ICM/TE marker expressions at morula stage, the increased expression of developmental genes, and other lineage markers for all three stages examined. Such ectopic expression likely contributes to the defective first and second cell fate specification at early and late blastocyst stages. It should also be noted that the lineage differentiation defects observed in this work is more severe than the maternal EED or EZH1/2 KO studies ^27, 60, 61^, which is likely due to depletion of both maternal and zygotic PRC2 by the dTAG approach.

In addition, our study also uncovered novel insights into bivalency establishment and illustrated how PRC2 regulates bivalency *in vivo*. By mapping H3K27me3 and H3K4me3 dynamics throughout early embryogenesis, we provide strong evidence indicating that bivalency acquisition mainly started in the E4.5 ICM and further increased in E6.5 EPI, which is in contrast to a previous study suggesting that bivalency is largely absent or infrequent in preimplantation embryos but established after implantation ^34^. Notably, during revision of our manuscript, a study reporting that bivalency is established at the E4.5 stage was published ^62^, which is consistent with our findings.

Previous studies have been mainly focused on how epigenetic factors directly regulate embryonic development ^37^. Our study on the role of PRC2 in PGC specification highlights the importance of lineage crosstalk, raising the possibility that epigenetic factors regulate embryogenesis via both cell-autonomous and non-cell-autonomous pathways. We showed that one of the notable direct PRC2 targets in E6.5 ExE is *Esrrb*, and it’s KO showed reduced *Bmp4* expression and PGCs numbers ^49^. Through a series of *in vivo* genetic manipulations and epigenome profiling, we demonstrated that PRC2 regulates *Bmp4* expression in ExE through the PRC2-ESRRB-BMP4 regulatory axis. Although our results highlight a role for PRC2-ESRRB-BMP4 signaling in promoting PGC expansion, BMP signaling during early germ cell specification is known to be spatially and temporally heterogeneous ^48^. In addition to BMP4 produced by the ExE, BMP ligands derived from the VE and extraembryonic mesoderm are also required for proper induction and patterning of PGCs ^63^. Although rapid PRC2 degradation has a pronounced effect on the ExE and only minimal effects on the EPI and VE, this does not exclude the possibility that PRC2 regulates PGC development through additional lineages and transcriptional programs beyond the ExE and ESRRB.

Acute PRC2 depletion at E6.5 embryos had minimal impact on the transcriptome in EPI but affected the expression of hundreds of genes in ExE. It remains unclear how acute loss of PRC2 causes different responses in these two lineages. It is possible that additional repressive chromatin markers that are specific to E6.5 EPI help maintain Polycomb repression in the absence of PRC2. This possibility raises a question: which epigenetic factors may interplay with PRC2 to silence bivalent genes in E6.5 EPI. PRC1 is one of the potential candidates, since it has also been proposed that H2AK119ub1 could be an important contributor to bivalency during preimplantation development ^44^. In addition, inducible PRC1 or PRC2 KO right after implantation suggest that PRC1 and PRC2 are largely independent from each other on the inactive X in extraembryonic tissues^64^. These observations suggest that PRC1/H2AK119ub1 and PRC2/H3K27me3 have different roles in regulating early embryogenesis. Further work is needed to systematically address how PRC1 along or together with PRC2 might regulate MZT, lineage specification, bivalency acquisition and gastrulation.

## Acknowledgements

We thank Drs. Qianying Yang, Yota Hagihara, Shan Jiang and Boyan Wang for their valuable discussions on the project and comments on the manuscript.

## Author contributions

Y.Z. conceived and supervised the project. C.Z. and Z.C. design the experiments. C.Z. performed all the experiments. M.W. performed all the bioinformatic analysis. Z.C., C.Z., M.W. and Y.Z. wrote the manuscript. All authors interpreted the data and reviewed the manuscript.

## Methods

### *Eed*^dTAG/dTAG^ mice generation

All experiments were conducted in accordance with the National Institute of Health Guide for Care and Use of Laboratory Animals and were approved by the Institutional Animal Care and Use Committee (IACUC) of Boston Children’s Hospital and Harvard Medical School (protocol number IS00000270-9). Mice were housed in specific pathogen-free facilities with regulated temperature (20-22°C) and humidity (40-70%), with a 12-hour light/dark cycle. To generate knock-in mice, the mixture of *Eed* donor DNA (30 ng/μl), sgRNA (40 ng/μl) and Cas9 mRNA (100 ng/μl) were injected into early 2-cell stage embryos with a Piezo-driven micromanipulator (Primer Tech, Ibaraki, Japan). Following injection, embryos were cultured in KSOM medium for 6 hours, and then transferred into the oviducts of pseudo-pregnant ICR female mice (Charles River). The F0 chimeras were backcrossed to C57BL/6J wild-type mice for a minimum of two generations to achieve stable genetic inheritance. The primers used for genotyping are listed in Supplementary Table 7.

### Zygotic KO embryo generation

For *Esrrb* and *Bmp4* zygotic KOs, the wide type or *Eed ^dTAG/dTAG^* background 1-cell embryos (6 hpf) were injected with the mixture of sgRNA (20 ng/μl) and Cas9 protein (20 ng/μl). After one day culture, 2-cell embryos were transferred into the oviducts of pseudo-pregnant ICR female mice (Charles River). Post-implantation embryos were collected at E6.5 or E7.0 for further analysis. Small piece of mixture of ExE and VE were used for genotyping. The zygotic *Eed* knockout was generated using similar methods, with sgRNAs designed according to a previous study ^37^. The sgRNAs and primers for genotyping are listed in Supplementary Table 7.

### Establishment of *Eed*^dTAG/dTAG^ cell line

*Eed*^dTAG/dTAG^ ESCs were generated by co-transfecting Eed-HAL-FKBPF36V-V5-HAR donor and px330 (Cas9 and sgRNA) plasmids into ES-E14TG2a cells (ATCC, CRL-1821) and cultured for 24 hours. Then transfected cells were subjected to puromycin selection (Gibco, A1113803) for 48 hours, followed by recovery in puromycin-free medium for 6-7 days. Single clones were picked for genotyping and further experiments. ESCs were maintained on 0.1% gelatin-coated plates under 2i/LIF culture conditions as previous used ^65^. The 2i/LIF culture medium consisted of DMEM (Gibco, 11960069) supplemented with 15% fetal bovine serum (Sigma-Aldrich, F6178), 1× MEM non-essential amino acids (Gibco, 11140050), 1 mM sodium pyruvate (Gibco, 11360), 100 U/mL penicillin-streptomycin (Gibco, 15140122), 0.084 mM β-mercaptoethanol (Gibco, 21985023), 2 mM GlutaMAX (Gibco, 35050061), 1000 IU/mL leukemia inhibitory factor (Millipore, #ESG1107), 0.5 μM PD0325901 (Tocris, 4192), and 3 μM CHIR99021 (Tocris, CHIR99021).

### *In vitro* fertilization and embryo culture

To induce superovulation, wild type C57BL/6J or *Eed*^dTAG/dTAG^ female mice (7-8 weeks old) were injected 7.5 IU of pregnant mare serum gonadotropin (PMSG, BioVendor, RP1782725000), followed by 7.5 IU of human chorionic gonadotropin (hCG, Sigma, C1063) 48 hours later. Oocyte-cumulus complexes (OCCs) were collected 14 hours after hCG injection. Before OCCs collection, sperm was obtained from the cauda epididymis of adult male mice (8-12 weeks old) and capacitated for 40 mins in 200 μl HTF medium (Millipore, MR-070-D). Then OCCs were collected and co-incubated with sperm for 6 hours in HTF medium. Zygotes with two-pronuclear were picked up and washed three times with warmed KSOM medium (Millipore, MR-106-D), then transferred to KSOM for further development at 37°C in a humidified 5% CO₂ atmosphere. All the culture medium was covered by mineral oil (Sigma, M5310-1L).

### Western blot

1.5×10^6^ cells were lysed in 200 µl 1× lysis buffer (50 μl 4 × LDS sample buffer, Invitrogen, NP0007; 20 μl 10 × reducing agent, Invitrogen, NP0007; 2 μl protease inhibitor, sigma, P8340; 128 μl H2O) and incubated on ice for 30 mins. Then the samples were heated at 98 °C for 10 mins, and transferred to ice immediately. 20 μl samples were loaded and run on NuPAGE 4-12% gel (Invitrogen, NP0322BOX), followed by transfer to 0.2 μm PVDF membranes. The primary antibodies anti-EED (1:500, CST, 85322S) and anti-β-actin (1:5000, CST, 4976S) were used for incubation at 4°C overnight. The membranes were washed for three times and incubated with Goat anti-Rabbit IgG (H+L) Superclonal™ Secondary Antibody-HRP (1:2000, Invitrogen, A27036) for 1 hour at room temperature. Target proteins were detected using ECL kit (Thermo Fisher Scientific, 32209) and imaged by iBright1500 (Invitrogen).

### Immunostaining and confocal microscope

The same Immunostaining protocol was used for both pre-implantation and post-implantation embryos. Briefly, embryos were fixed in 4% paraformaldehyde/0.5% triton for 20 mins. Following fixation, embryos were washed three times in PBS containing 0.1% Triton X-100 and subsequently blocked in PBS supplemented with 1% bovine serum albumin (BSA) and 0.1% Triton X-100 for 1 h at room temperature. Primary antibody incubations were performed overnight at 4°C using the following dilutions in blocking buffer: anti-EED (1:500, CST, 85322S), anti-H3K27me3 (1:200, Active Motif, 61017), anti-V5 (1:200, Invitrogen, R960-25), anti-OCT4 (1:200, Santa Cruz, sc-5279), anti-GATA4 (1:200, R&D Systems, MAB2606-SP), anti-NANOG (1:200, Abcam, ab80892), anti-CDX2 (1:500, R&D Systems, AF3665-SP), anti-SOX2 (1:200, R&D Systems, AF2018-SP) and anti-TFAP2C (1:200, Proteintech, 14572-1-AP). After three times of washing, the embryos were incubated with secondary antibodies for 1 hour at room temperature: Donkey anti-Mouse IgG-Alexa Fluor 568 (1:500, Invitrogen, A10037), Donkey anti Rabbit IgG (H+L)-Alexa Fluor 488 (1:500, Invitrogen, A-21206) and Donkey anti-Goat IgG (H+L)-Alexa Fluor 647 (1:500, Invitrogen, A-21447). Hoechst 33342 (10 μg/ml, Sigma) was used for DNA staining. Embryos were imaged with a confocal microscope (Zeiss, LSM800).

Two complementary approaches were used for the cell counts shown in Figs. 1j-k, depending on nuclear density and morphology. For the majority of embryos, in which individual nuclear and lineage markers could be readily distinguished, total cell numbers and lineage identities were determined by direct manual counting under the microscope, using lineage-specific fluorescence channels. For embryos in which the ICMs appear more tightly compacted, full 3D z-stack imaging was performed to ensure accurate counting. Embryos were imaged through their entire volume with a z-step size of 2 µm, and nuclei were identified and counted by manual inspection of sequential optical sections using ImageJ, based on DNA staining. Lineage identity was assigned using CDX2 or OCT4, evaluated consistently across the full z-stack.

### Embryo isolation

For E4.5 ICM isolation, E4.5 embryos were treated by Acidic Tyrode’s solution (Millipore) to remove the zona pellucida. Embryos were first incubated with anti-mouse serum antibody (1:3 dilution in KSOM medium, Sigma-Aldrich, M5774) for 30 min at 37°C, followed by guinea pig complement (1:3 dilution in KSOM, Millipore, S1639) treatment for 30 min at 37°C. The ICMs were isolated by removing trophectoderm cells with a glass pipette. E4.5 ICM refers to the mixture of EPI and PrE cells. Due to technical difficulties, EPI and PrE cannot be separated. For E6.5 embryo isolation, embryos were dissected from the decidua, followed by removal of the Reichert’s membrane. Embryos were then incubated in 100 μl 0.25% trypsin supplemented with 2.5% pancreatin (Sigma, P3292) on ice for 15 minutes. The enzymatic digestion was quenched by adding 100 μl of 10% FBS. The VE layer was manually removed using a glass pipette, the EPI and ExE were isolated using a sharp needle under a dissection microscope.

### dTAG treatment

To degrade EED in pre-implantation embryos, *Eed*^dTAG/dTAG^ embryos were cultured in KSOM with 1 μM dTAG13 (Tocris, 6605). To degrade EED in ESCs, *Eed*^dTAG/dTAG^ ESCs were cultured in 2i/LIF medium with 0.5 μM dTAG13. For EED degradation in post-implantation embryos *in vitro*, *Eed*^dTAG/dTAG^ embryos were maintained in culture medium (75% pregnant rat serum, 2 mM GlutaMAX, 11 mM HEPES and 1x MEM NEAA) containing 2 μM dTAG^V^-1. For EED degradation in post-implantation embryos *in vivo*, intraperitoneal (IP) injections were performed. Specifically, 1 mg dTAG^V^-1 was initially dissolved in 25 μl DMSO and subsequently diluted with 475 μl 10% castor oil (Sigma, C5135) to prepare the injection solution. Mice (∼28 g each) were injected with 1 mg dTAG^V^-1 every 12 hours to maintain sustained EED degradation.

### CUT&RUN, ATAC-seq and RNA-seq libraries preparation and sequencing

CUT&RUN was performed as previously described ^65^. Briefly, intact or isolated embryos were mixed with activated Concanavalin A Magnetic Beads (Polysciences, 86057-3) for 10 mins at RT, followed by incubation with primary antibodies overnight at 4°C. After three washes, samples were included with 3 ng/μl pA-MNase (home-made) for 2 hours with rotation at 4 °C. Subsequently, pA-MNase was activated by incubating with 200 μl pre-cooled 0.5 μM CaCl2 for 20 mins at 4 °C, and quench by adding 23 μl 10×stop buffer. To release DNA, samples were first incubated at 37 °C for 15 minutes, followed by addition 2.5 μl 10% SDS and 2.5 μl Proteinase K (20 mg/ml; Thermo Fisher, AM2546), and then incubated at 55 °C for 1 hour. DNA was subsequently purified using phenol-chloroform extraction. DNA library was constructed using NEBNext Ultra II DNA library preparation kit (New England Biolabs, E7645S). Primary antibodies anti-H3K4me3 (1:100, CST, 9727S), anti-H3K27me3 (1:100, Diagenode, C15410069), anti-H2AK119ub1 (1:100, CST, 8240S), and anti-ESRRB (1:100, Proteintech, 22644-1-AP) were used for CUT&RUN.

ATAC-seq was performed as previously described^66^. Briefly, isolated embryos were incubated with Tn5 (Diagenode, C01070012-30) for 15 mins at 37°C. Subsequently, DNA fragments were released by dissecting with Proteinase overnight at 55°C. DNA libraries were constructed with NEBNext High-Fidelity 2×PCR Master Mix (NEB, M0541S).

For RNA-seq, intact or isolated embryos were collected and immediately frozen at -80 °C until use. SMART-Seq™ v4 Kit (Clontech, 634890) was used to construct cDNA libraries, and Nextera® XT DNA Sample Preparation Kit kit (Illumina, FC-131-1024) was used for the sequence libraries construction.

All libraries were sequenced by the Illumina NextSeq 1000/2000 system.

### RNA-seq data analysis

The raw paired-end sequencing reads of RNA-seq were first processed with the Trimmomatic (v0.39)^67^ to remove sequencing adaptors if present. After adaptor trimming, reads with length less than 35 bp were discarded. The trimmed reads were mapped to GRCm38 mouse reference genome using STAR (v2.7.8a)^68^. The gene annotations were downloaded from GENCODE (https://www.gencodegenes.org) with version vM24. The read counts mapped to each gene were calculated using RSEM (v1.3.1)^69^ with the transcriptome alignments generated by STAR as input. To calculate differentially expressed genes (DEGs) between conditions, R package DESeq2 (v1.46.0)^70^ was used. Genes with adjusted *p*-value < 0.05, fold change ≥ 2 and mean FPKM ≥ 1 were identified as DEGs. The Gene Ontology (GO) enrichment of DEGs was calculated with R package clusterProfiler (v4.14.3)^71^.

### CUT&RUN data analysis

The raw paired-end reads of CUT&RUN were trimmed with Trimmomatic (v0.39)^67^ to remove sequencing adaptors and the trimmed reads with length at least 35 bp were kept. The cleaned reads were mapped to GRCm38 mouse reference genome using bowtie2 (v2.4.2)^72^ with parameters: --local --very-sensitive-local --no-unal --no-mixed --no-discordant --dovetail -I 10 -X 700 --soft-clipped-unmapped-tlen. Picard MarkDuplicates (v2.23.4) were used to remove PCR duplicates. The aligned reads were further filtered to retain proper paired reads with a minimum mapping quality of 30. For histone modifications CUT&RUN data, the FPKM (fragments per kilobase region per million mapped fragments) signal tracks were generated with bamCoverage in deeptools (v3.5.1)^73^ with 100 bp bin size. For TF CUT&RUN data, the CPM (counts per million) signal tracks were generated with bamCoverage in deeptools (v3.5.1) with 1 bp bin size. To call peaks for H3K4me3 ChIP-seq data and ESRRB CUT&RUN data, the MACS2 (v2.2.7.1)^74^ was used with parameters: -f BAMPE -B --SPMR -p 1e-4 -g mm. The reproducible peaks between two replicates were calculated with the irreproducible discovery rate (IDR) framework^75^, with the IDR cutoff of 0.05. For ESRRB binding peaks, we further filtered them with q-value cutoff of 10^−30^ to remove weak binding peaks. The heatmaps of CUT&RUN signals around TSSs or peak centers were calculated with computeMatrix in deeptools (v3.5.1) with bin sizes of 10, and plotted with R packages profileplyr (v1.22.0) and EnrichedHeatmap (v1.36.0). The promoter (±1 kb around TSS) average levels of H3K27me3 or H3K4me3 were calculated with multiBigwigSummary in deeptools (v3.5.1). For genes with multiple TSSs, the highest value among all the TSSs was used as the signal level of that gene.

### ATAC-seq data analysis

The signal tracks of ATAC–seq were calculated with the ENCODE ATAC–seq pipeline using default parameters (v2.1.2, https://github.com/ENCODE-DCC/atac-seq-pipeline), and the fold change signal tracks were used.

### Histone modification signal normalization across stages

To study the H3K27me3 or H3K4me3 levels across different developmental stages, the FPKM values need to be normalized to make them comparable. We used a normalization method as previously described^76, 77^. Briefly, we first calculated the average FPKM at gene promoter regions (-1,000 bp to +500 bp around TSS) and ranked them in descending order. For each stage, the median FPKM value of the top 3,000 promoters were considered as the saturated signal value of that stage. Then the saturated value of each stage was divided by the saturated value of ESC (or ICM) stage to get the scale factor for that stage. Finally for each stage, the original FPKM values were multiplied by the scale factor of that stage to obtain the normalized FPKM.

### Bivalent gene identification

Bivalent genes were identified based on the normalized FPKM of H3K27me3 and H3K4me3 at promoter regions. For all the stages, genes with both signal values above a unified threshold were considered as bivalent genes. To find the threshold, we used the number of overlapped peaks between H3K4me3 and H3K27me3 in E4.5 ICM as a reference. Specifically, the H3K4me3 peaks in E4.5 ICM were called as described above. The average raw FPKM of H3K27me3 at these regions (±2 kb around H3K4me3 peak center) was calculated and regions with a value ≥ 3 were considered as strong H3K27me3 domains overlapping with H3K4me3 peaks, which resulted in 1,457 such bivalent regions in E4.5 ICM. Then we searched a threshold value θ (θ = 6.365) so that when the genes promoter normalized FPKM of H3K27me3 ≥ θ and the normalized FPKM of H3K4me3 ≥ θ in E4.5 ICM, the same number (1,457) of regions could be derived. Finally, for each stage, genes with promoter normalized FPKM of H3K27me3 ≥ 6.365 and the normalized FPKM of H3K4me3 ≥ 6.365 were identified as bivalent genes of that stage.

### Public data sets used

RNA-seq of mouse pre-implantation embryos^78, 79^: GSE71434 and GSE76505. Ribo-seq of mouse pre-implantation embryos^35^: GSE169632. H3K27me3 ChIP-seq of mouse pre-implantation embryos^45^: GSE73952. H3K4me3 ChIP-seq of mouse pre-implantation embryos^78^: GSE71434. H3K4me3 ChIP-seq of mouse post-implantation embryos^80, 81^: GSE125318 and GSE124212. TFAP2C and SOX2 CUT&RUN of E6.5 ExE embryos^82^: GSE216256.

### Statistics and reproducibility

Student’s *t*-tests were performed for graph analysis. All the other statistical analyses methods were showed in the individual figure legends. No statistical methods were used to pre-determine sample sizes, but our sample sizes are similar to or greater than those reported in previous publications. No data were excluded from the analyses. The experiments were not randomized. The investigators were not blinded to allocation during experiments and outcome assessment.

## Data and materials availability

All data generated in this study have been deposited to the NCBI Gene Expression Omnibus (GEO) with accession number GSE298615.

---

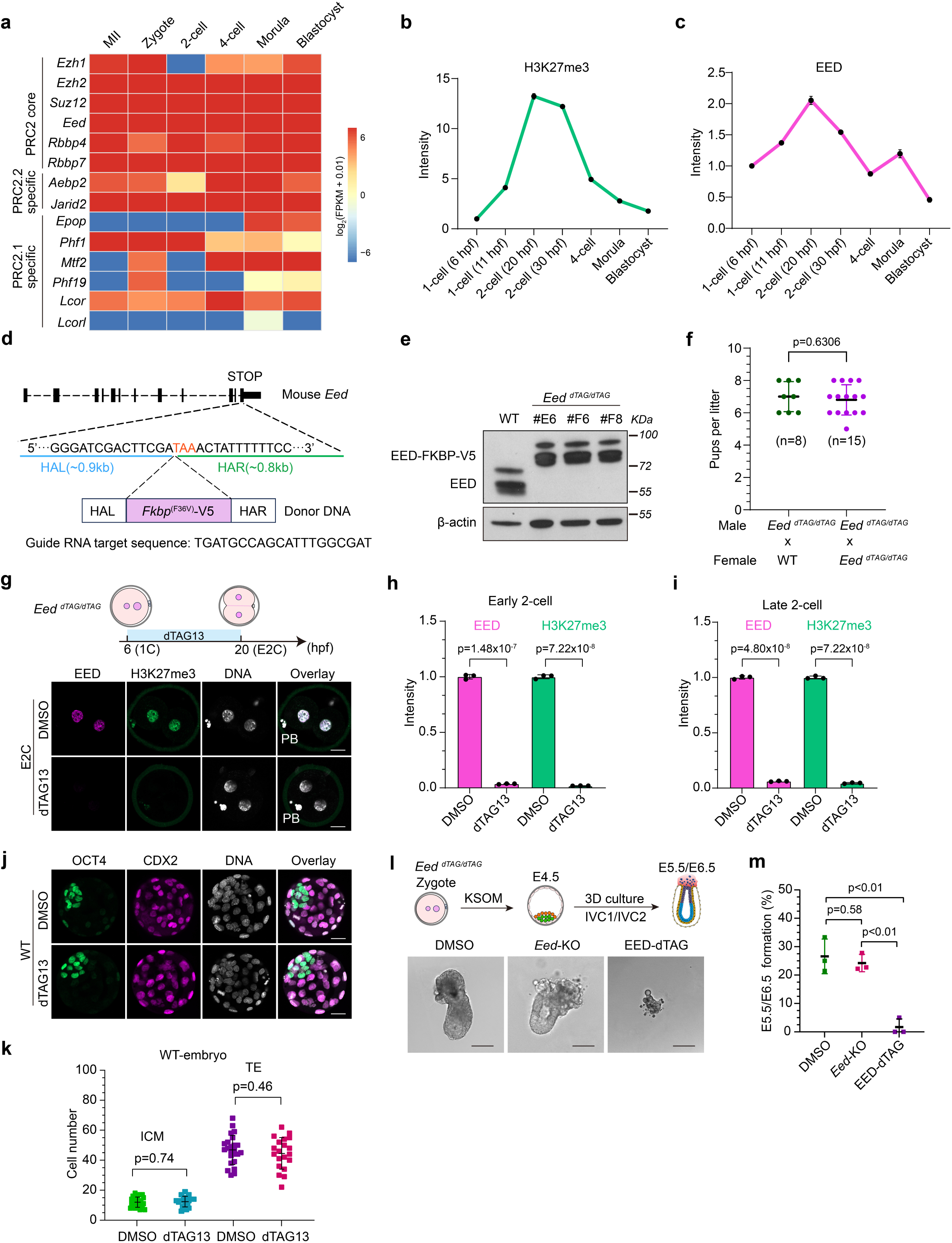

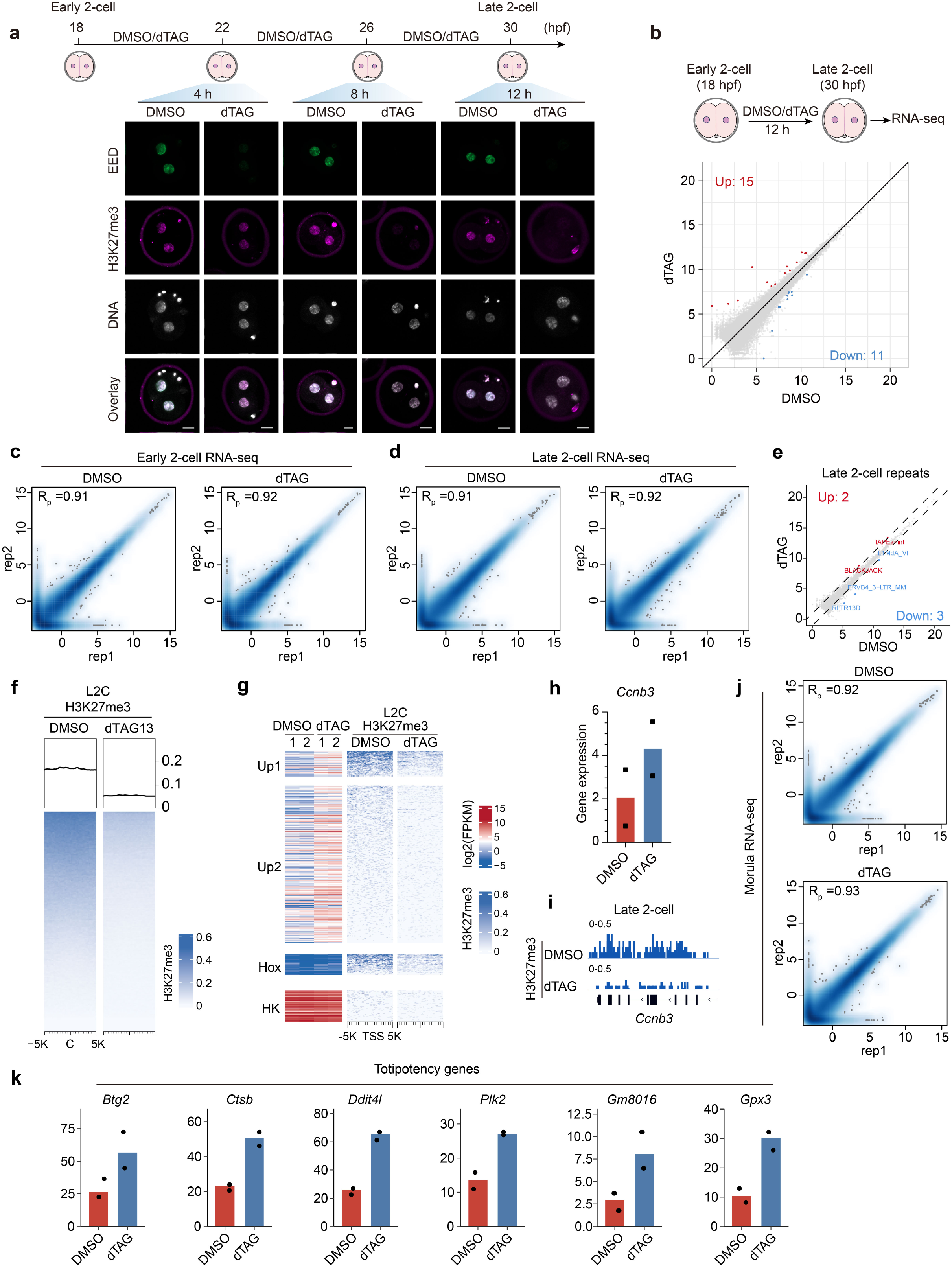

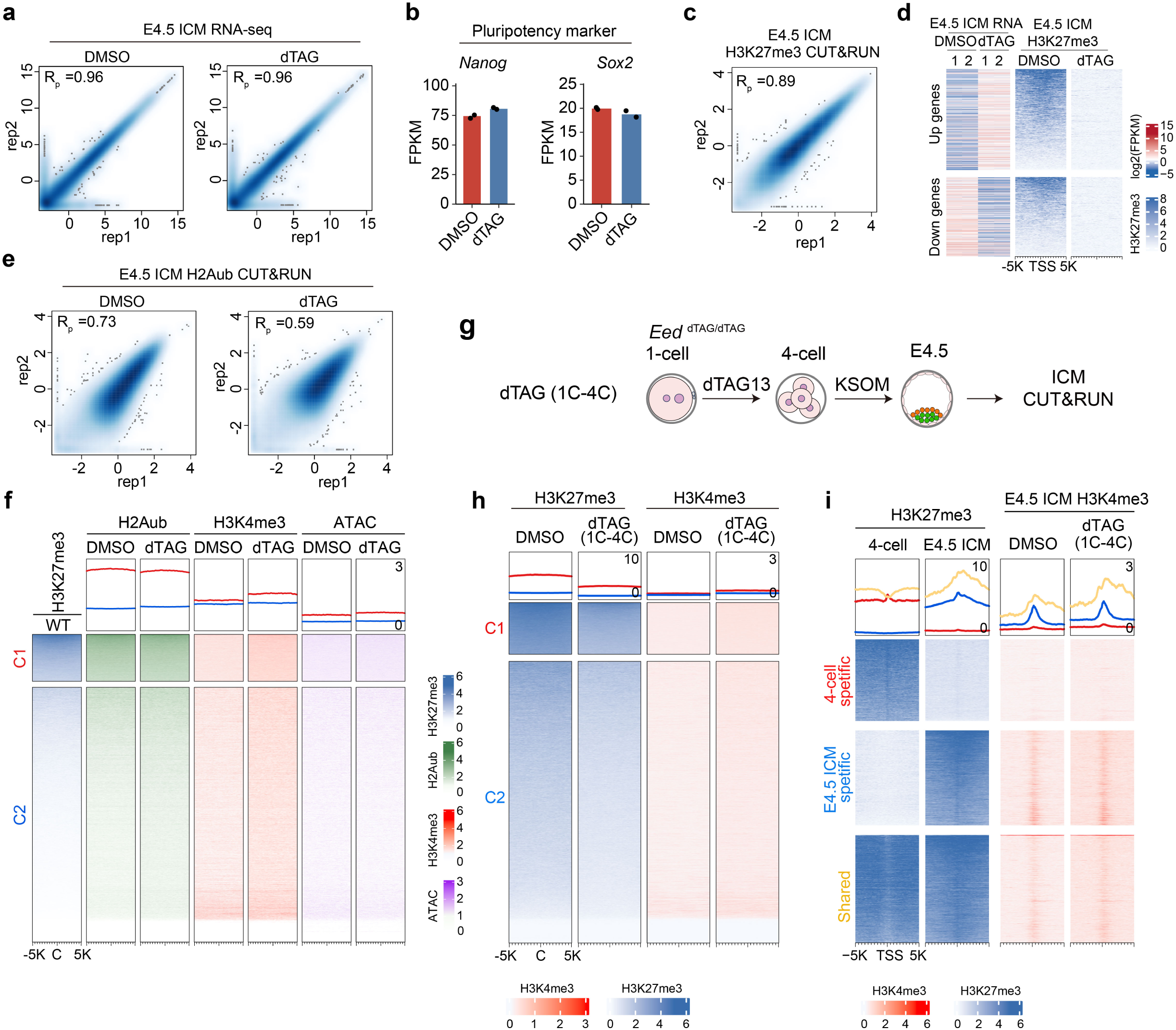

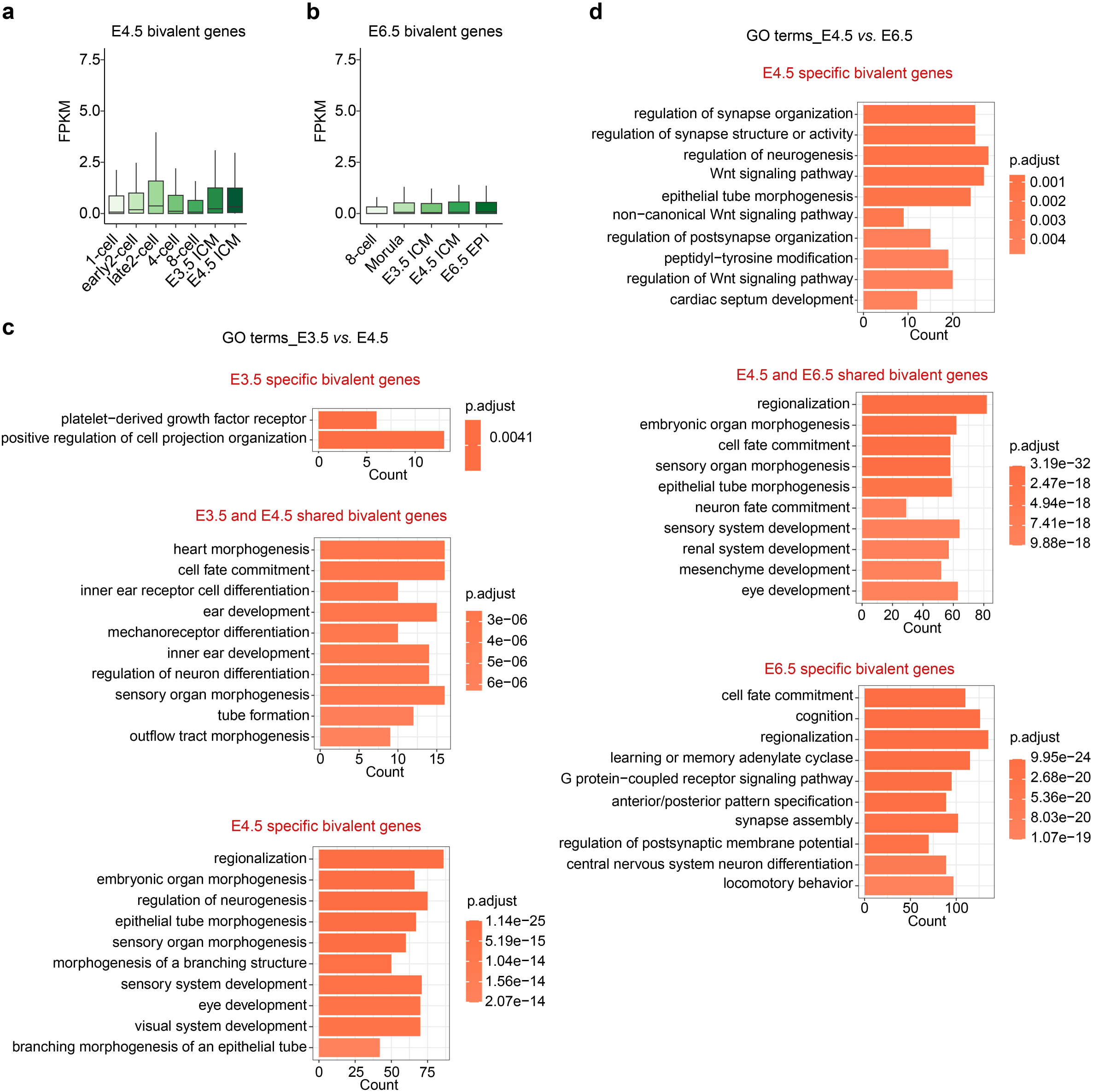

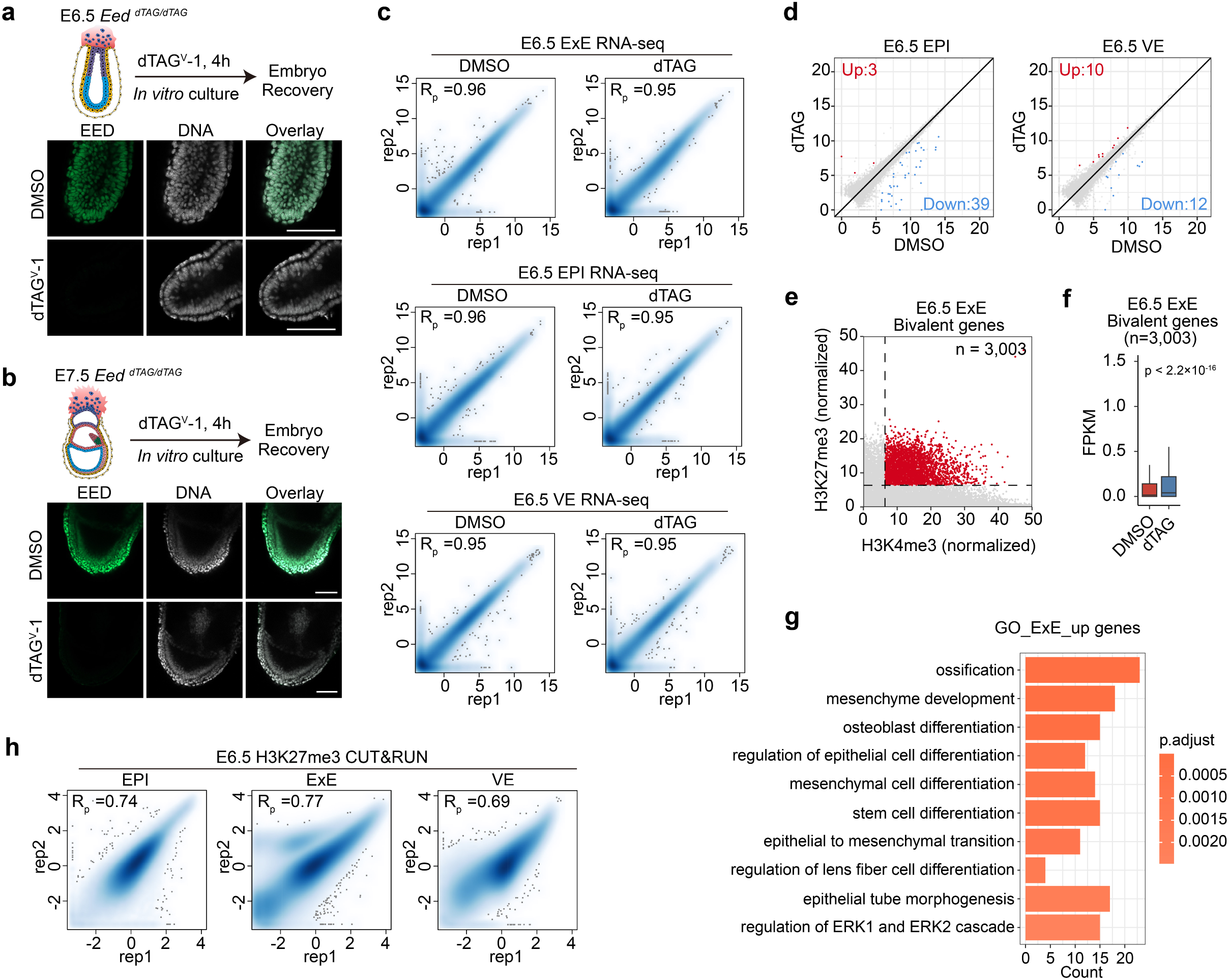

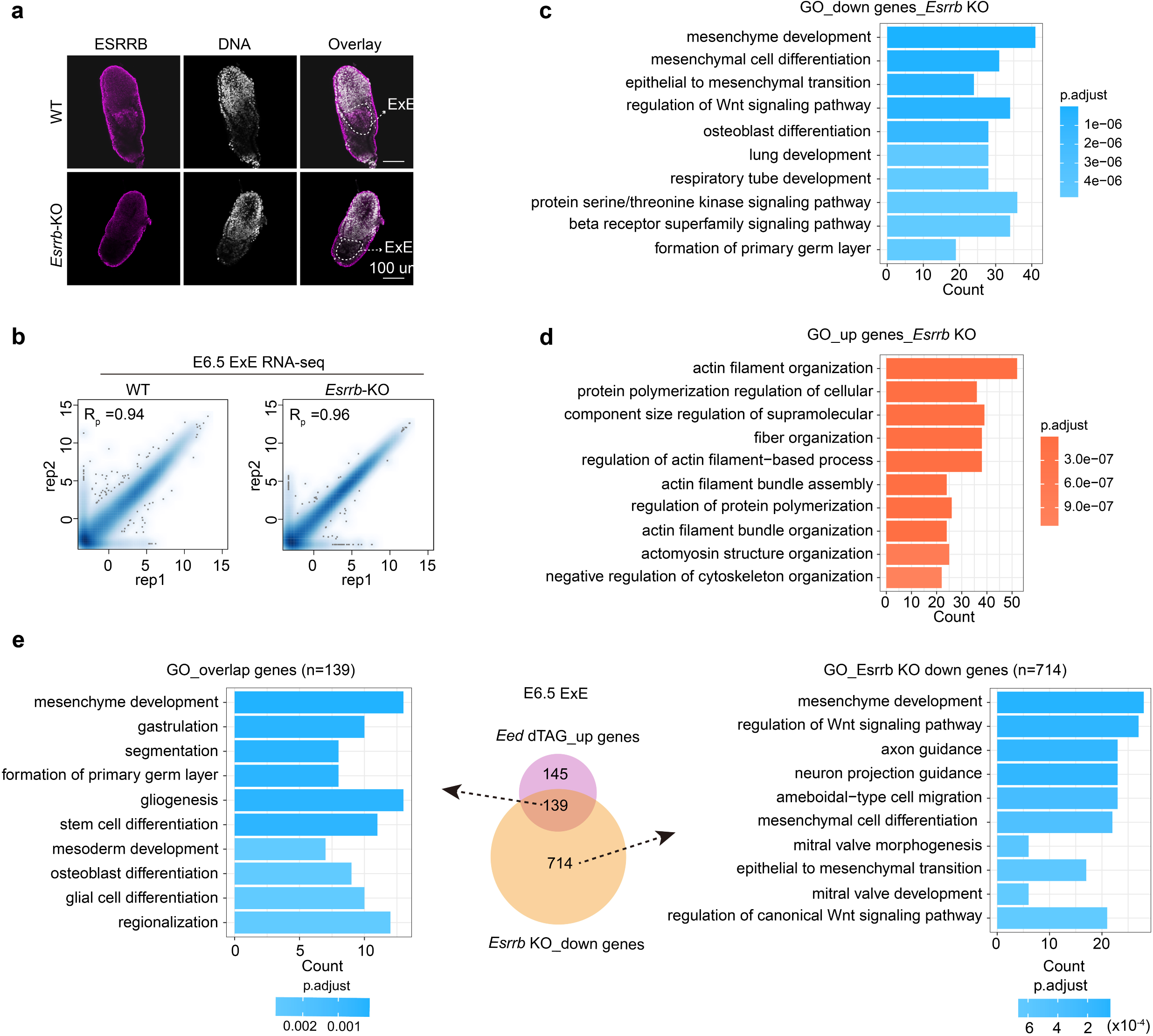

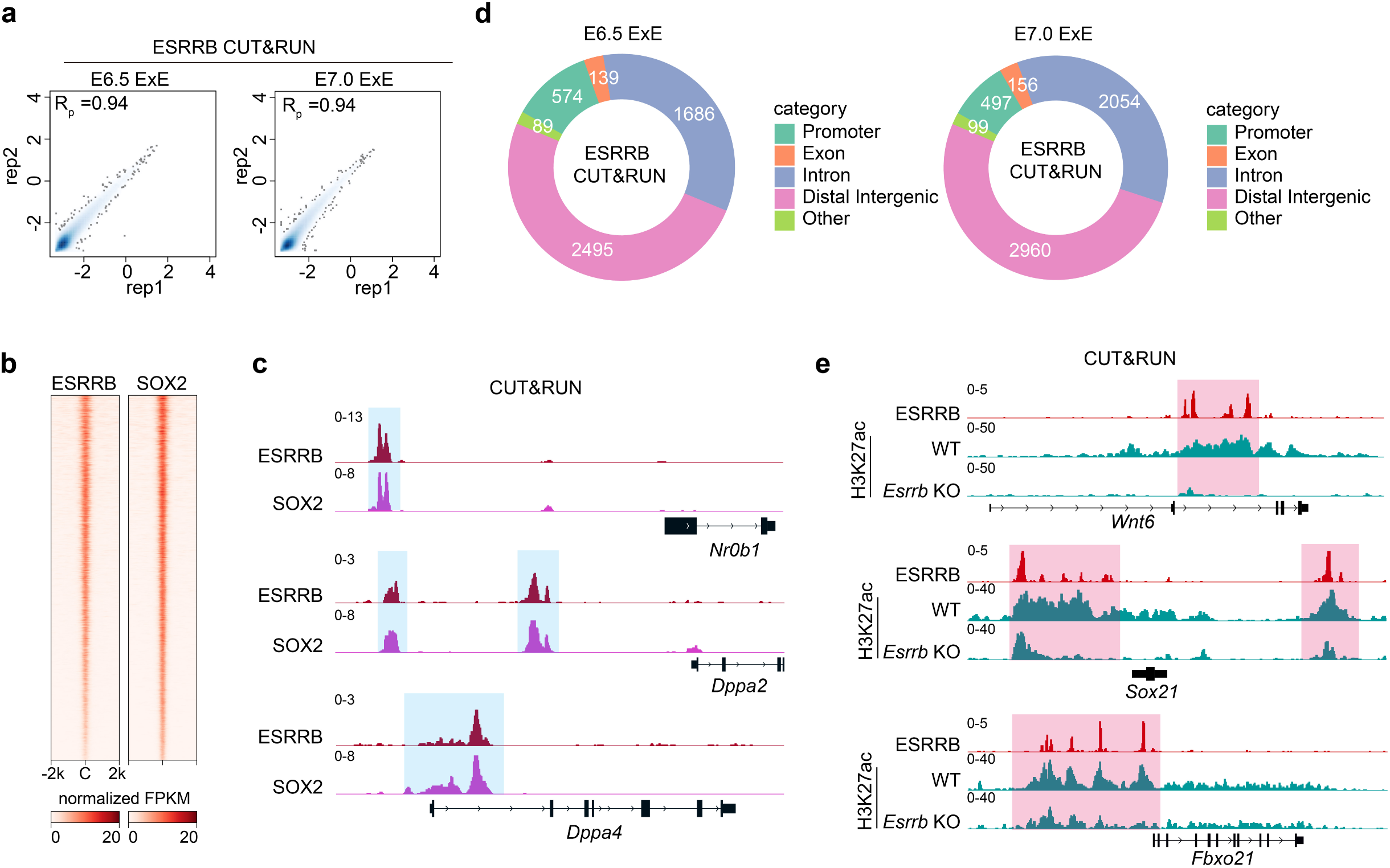

